# A Design-of-Experiments Strategy for Engineering 3D Topographical Features in Osteosarcoma Modelling

**DOI:** 10.1101/2025.09.03.674028

**Authors:** Kozim Midkhatov, George Taylor, Lee Stevens, Mahetab H. Amer

## Abstract

Osteosarcoma is a highly aggressive bone cancer with poor patient outcomes, partly due to the limited predictive power of current preclinical models. Conventional two-dimensional (2D) cultures fail to recapitulate physiologically relevant cell-matrix interactions, while animal models suffer from inter-species variability. To investigate how extracellular matrix (ECM) topographical features influence osteosarcoma behaviour, polylactic acid-based microparticles were engineered with bone-mimetic stiffness and defined surface topographies, guided by a Design-of-Experiments (DoE) approach. This enabled systematic variation of microparticle architecture (8-63 µm diameter; 2-13 µm dimple size) for studying the impact of surface topography on osteosarcoma cell behaviour in 3D culture, with doxorubicin treatment as a functional test to evaluate the effect of 3D topographical cues on chemotherapy sensitivity. Topography modulated cell-microparticle aggregation dynamics. At 96 hours post-seeding, MG-63 cells displayed significantly reduced metabolic activity on all 3D microparticle designs, with heterogeneously dimpled-topography cultures displaying significantly lower DNA content than conventional 2D cultures. In U2OS cells, metabolic activity was significantly lower on smooth microparticles compared to dimpled designs, with all 3D cultures showing significantly lower DNA content versus 2D. Response to doxorubicin was more strongly influenced by culture dimensionality than surface topography, underscoring the importance of 3D context. Significant metabolic differences between 3D and 2D cultures were observed, including the enrichment of amino acid related pathways and downregulation of ferroptosis signatures in 3D microparticle cultures. Topography displayed subtler effects on lipid and nucleotide metabolism. This study highlights how topographically-patterned 3D substrates can shape osteosarcoma cell behaviour and drug response for disease modelling applications. Our DoE-guided platform enables systematic investigation for dissecting how ECM-inspired physical cues influence osteosarcoma progression and therapeutic resistance.

**Graphical Abstract:** 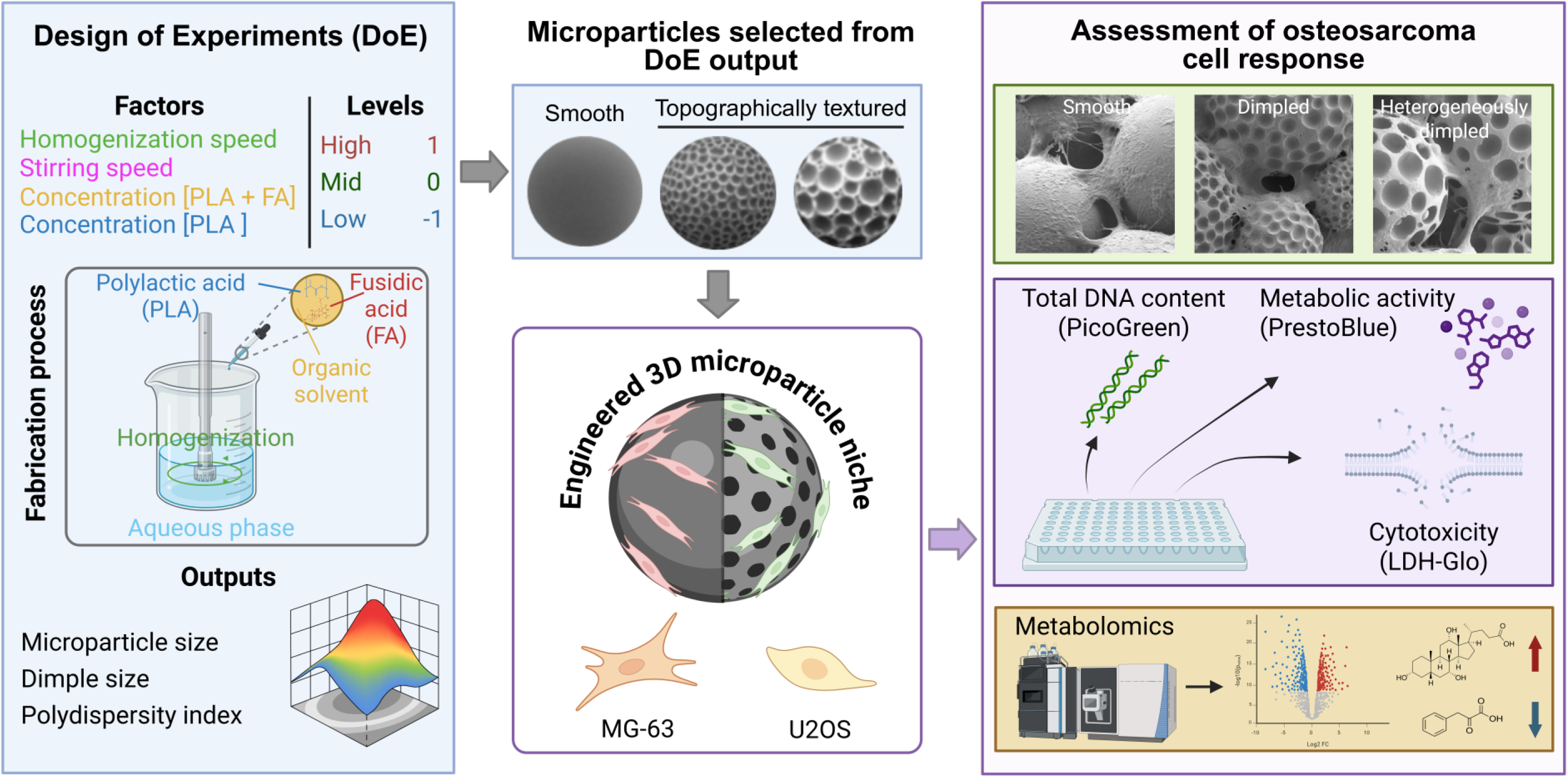

## 1. Introduction

Osteosarcoma, the most common primary bone malignancy, poses significant clinical challenges due to its aggressive nature, high metastatic potential, and resistance to conventional therapies [1]. Despite advances in surgical and chemotherapeutic strategies, prognosis for metastatic or recurrent disease remains poor [1]. One of the key barriers to therapeutic discovery is the lack of preclinical models that faithfully recapitulate the tumour microenvironment (TME), as conventional 2D *in vitro* models are devoid of proper cell-extracellular matrix (ECM) interactions, while animal models suffer from interspecies variation [2]. Moreover, challenges that limit the discovery and progression of new therapies include tumour heterogeneity in osteosarcoma, and limited targetability due to poorly defined tumour antigens [3]. Failure of candidate therapies in clinical trials, despite showing promise in the laboratory, highlights the translational gap in current drug development pipelines. For example, trabectedin showed promise against drug-resistant osteosarcoma cell lines in vitro [4], but failed to demonstrate clinical efficacy in phase II clinical studies [5]. Similarly, robatumumab resulted in tumour shrinkage in xenograft models [6, 7] while failing to demonstrate clinical efficacy in patients with metastatic osteosarcoma [8]. Such discrepancies highlight the limitations of current 2D cultures and animal models, which fail to capture the spatial, mechanical, and biochemical complexity of the osteosarcoma microenvironment.

The ECM has been recognized as a critical factor influencing tumour progression and resistance to therapies [9]. In recent years, three-dimensional (3D) tissue-engineered cancer models have emerged as an improved representation of tumour biology and drug response compared to traditional 2D cultures [10]. However, materials commonly employed for osteosarcoma modelling, such as soft hydrogels (1–150 kPa) fabricated using natural or synthetic polymers [11], present notable limitations. The animal-derived Matrigel suffers from batch-to-batch variability [12, 13] and weak mechanical properties [14], while collagen is often combined with other materials for structural support [15]. Moreover, alginate [16], gelatin [17] and polyethylene glycol [18], in their typical formulations used for osteosarcoma modelling, exhibit low mechanical stiffness and are mechanically weak, limiting their capacity to mimic the native bone environment [19, 20]. Bone is a mechanically stiff tissue (GPa scale) [21]. Malignant osteoblastic cells also deposit disorganised osteoid matrix [22], and therefore it is essential for in vitro models to capture both the mechanical and architectural features of the TME. Current in vitro models rarely replicate these defining features, limiting their utility for investigating osteosarcoma-specific cell-matrix interactions.

Several studies have attempted to replicate the complexity of the osteosarcoma microenvironment using hydrogel-or polymer-based systems incorporating osteoconductive components, such as microribbon scaffolds containing gelatin and hydroxyapatite [23], polylactic acid (PLA)-based scaffolds [24, 25], and polyurethane constructs [26]. While these approaches offer some compositional relevance, they often lack the microscale physical features that modulate tumour cell behaviour and may not be amenable to high-throughput systematic interrogation of cell–matrix interactions. Recent studies have begun to study how osteosarcoma cells dynamically interact with their ECM, revealing new opportunities to interrogate disease mechanisms [27]. Lu *et al.* developed an osteosarcoma-on-a-chip system using decellularized tumour-derived ECM, enabling matrix-mediated control of drug responses [28]. Yet, the ECM also presents spatially patterned 3D topographies that are increasingly recognised as important regulators of cell behaviour [29]. Importantly, the disruption of normal topographical cues may contribute to pathological processes, including cancer progression [30]. Studies have demonstrated that specific topographical patterns can promote skeletal stem cell differentiation [31] and modulate osteosarcoma cell responses [32]. Recent work has shown that cancer cells can respond differentially to topographical features compared to their normal counterparts, with altered migration patterns, and invasion capacity observed on patterned 2D surfaces [33, 34]. Collectively, these advances highlight the need for biomimetic model systems that can capture key matrix parameters in vitro. However, the role of 3D topographical cues in osteosarcoma cell behaviour remains largely unexplored.

Here, we present a platform based on topographically-patterned PLA microparticles designed to mimic bone-relevant stiffness (with Young’s modulus >1 GPa) and microscale topography [35]. We have previously shown that these topographically textured microparticles induce osteogenesis of mesenchymal stem cells (MSCs) without the addition of osteoinductive supplements [35, 36]. In this study, we extend this approach by applying a Design of Experiments (DoE) strategy to systematically examine the influence of defined 3D surface features on osteosarcoma cell response. By incorporating physiologically relevant matrix stiffness with engineered 3D topographical cues, this platform offers a tuneable and scalable system for dissecting osteosarcoma cell-matrix interactions and supporting therapeutic development.

## 2. Materials and methods

### 2.1 Fabrication of microparticles

Polylactic acid (PLA) microparticles were fabricated by a solvent evaporation oil-in-water emulsion technique, as previously described [35]. The organic phase consisted of PLA (Viatel DL 09 E, IV=0.87 dL/g; Ashland) and fusidic acid (FA) (#458460050, ThermoFisher Scientific) in dichloromethane (DCM) as solvent (L090000, Sigma-Aldrich). This was homogenized in an aqueous phase consisting of 1% w/v of poly(vinyl alcohol) (PVA) (348406, Sigma-Aldrich) at a set speed with a homogenizer (Silverson L5M, Silverson Machines). The resulting emulsion was then stirred at room temperature to allow solvent evaporation. Microparticles were then washed with phosphate buffered saline (PBS) (12821680, ThermoFisher Scientific), after which they were filtered through 100 μm cell strainers to remove polymer films, debris, and fabrication artefacts, then freeze dried and characterised. Only the <100 μm filtered fraction was used in subsequent experiments and DoE analyses across all runs.

### 2.2 Design of Experiments (DoE)

A DoE approach was implemented to efficiently investigate the effects of multiple fabrication parameters on microparticle design characteristics. A central composite design (CCD) was chosen as part of a response surface methodology (RSM) for the optimisation of the fabrication process. A three-level composite design was used to screen the effects of the four selected factors: homogenization speed, total mass concentration of PLA and fusidic acid in the solvent (concentration [PLA+FA]), concentration of PLA within the PLA-fusidic acid mixture (concentration [PLA]) and stirring speed (Table 1). Twenty-six unique conditions were each run in duplicate (52 runs in total), and all runs were performed in randomized order to minimize systematic biases. Each experimental run produced a batch of microparticles, which were characterized for the following output variables: (a) microparticle mean size (diameter), (b) surface dimple (topography feature) mean size, and (c) polydispersity index (PDI) of size distribution for both the microparticle size and the dimples. These response metrics were chosen to quantify the key attributes of particle size and surface texture uniformity.

**Table 1.** Fabrication factors and corresponding levels used in the central composite design model

| **Fabrication factors (units)** | **Level I (-1)** | **Level II (0)** | **Level III (+1)** |
| --- | --- | --- | --- |
| **Homogenization speed (rpm)** | 1800 | 3000 | 4200 |
| **Stirring speed (rpm)** | 180 | 390 | 600 |
| **Concentration [PLA+FA] (%)** | 5 | 12.5 | 20 |
| **Concentration [FA] (%)** | 20 | 50 | 80 |

Statistical analysis of DoE results was carried out using the built-in RSM modelling tools in JMP Pro 18 (SAS Institute, USA). Analysis of variance (ANOVA) was used to assess the significance and relative influence of each fabrication factor on the measured outcomes, including key two-factor interactions affecting microparticle size and surface topography.

Model terms with a *p*-value < 0.05 were considered statistically significant. Model performance was evaluated using diagnostic metrics calculated in JMP Pro 18. The quality of fit for each response surface model was assessed using the coefficient of determination (R²) and predicted R² (Q²). Response surface plots were generated to visualize how each factor influenced the responses, facilitating the determination of optimal factor settings.

The model’s predictive performance was evaluated by fabricating five additional validation experimental runs of random combinations within the design space (but not used in building the model). Results from these validation runs were compared against the model-predicted values for microparticle and dimple sizes. The prediction errors were used to calculate the predictive R² (Q²), a measure of the model’s predictive accuracy. A high predictive Q² (close to 1.0) would confirm the robustness of the CCD-derived model to reliably predict microparticle design attributes for untested fabrication conditions. Q² was calculated using the mean response of the validation set as the reference with the following formula:

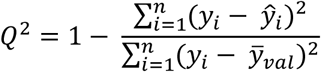

where 𝑦_𝑖_ – actual values from validation set, 𝑦^_𝑖_ – values predicted by the model, 𝑦̅_𝑣𝑎𝑙_ – mean of the actual values in the validation set, and n – total number of samples in the validation set.

### 2.3 Microparticle size analysis

Microparticles were dispersed in deionised water, and an adequate volume of this suspension was added to a beaker of deionized water until a laser obscuration value of at least 4.5% was obtained. Size distribution was determined using a Malvern MasterSizer 3000 (Malvern Panalytical, United Kingdom) equipped with the HydroEV dispersion unit. Mastersizer Xplorer software (v5.20) was used for analysis. PDI was used as a measure of broadness of size distribution.

### 2.4 Scanning Electron Microscopy (SEM)

Microparticles were imaged on a scanning electron microscope (TESCAN VEGA3) by first sputter-coating the samples with a thin layer (7nm) of gold/palladium (Q150T ES Plus, Quorum Technologies), then imaging at 5 kV. In the case of cell-containing aggregates, samples were sequentially dehydrated with ethanol before SEM imaging (TESCAN Mira3 SC). They were coated with gold/palladium for 5 minutes and imaged at 2kV. Dimple size was characterised using ImageJ software by measuring the diameter of a minimum of 250 dimpled microparticles over 5 individual SEM images for two fabricated batches.

### 2.5 Atomic force microscopy (AFM)

To map the surfaces of the different microparticle designs, AFM was performed using a BioScope Resolve (Bruker Corporation, United States) mounted on an Eclipse Ti-U (Nikon) inverted optical microscope, and AFM probe ScanAsyst-Air (Bruker Nano Inc; 0.4 N/m). Imaging was performed using ScanAsyst Peak-Force Tapping mode in air with a scan size of 15 μm at a scan rate of 0.799-1 Hz. The instrument is regularly calibrated using a grating with 180 nm deep and 10 µm x 10 µm depressions. Images were processed by applying flattening of second order using Nanoscope Analysis software (v3.0).

### 2.6 BET surface area measurements

Specific surface area of the microparticles was determined using the Brunauer–Emmett–Teller (BET) method applied to Krypton sorption isotherms using an ASAP 2420 instrument (Micromeritics Corporation, United States), as described before [36]. Prior to measurement, samples (250 mg) were degassed at 37 °C under high vacuum (<0.013 mbar) for 48 hours to remove adsorbed moisture and other gases. Krypton sorption isotherms were recorded at - 196°C over a relative pressure (P/P0) range of 0.00 to 0.65. BET surface area was calculated from the linear portion of the BET plot, between a P/P0 range of 0.05 to 0.30 ensuring a positive BET constant and monolayer capacity within the selected pressure range using Microactive Software V7.0. Pore volume and size distribution (1.75–20.00 nm) were determined from the adsorption isotherm using the Derjaguin–Broekhoff–de Boer model. The cross-sectional area of krypton was taken as 0.210 nm². Micropore volume was calculated at 2 nm, and limited mesopore volume from 2-20 nm using pore size vs cumulative pore volume using the Derjaguin–Broekhoff–de Boer model. Where additional batches of microparticles were required, they were fabricated under the same conditions as the original BET-characterised samples, and consistency in microparticle and dimple size was confirmed using SEM and laser diffraction size analysis.

### 2.7 Osteosarcoma cell culture

Two established osteosarcoma cell lines, MG-63 (cat. no. 86051601, ECACC) and U2OS (cat. no. 92022711, ECACC) were cultured in Dulbecco’s Modified Eagle Medium (DMEM; 41965, Gibco) and McCoy’s 5A (M9309, Merck) culture media, respectively. Medium was supplemented with 10% FBS (F9665, Sigma-Aldrich), 1% Penicillin/Streptomycin and 1% L-Glutamine (Gibco).

### 2.8 Preparation of microparticles for cell culture studies

Selected microparticle designs (Table 1) were placed in CELLSTAR^®^ 96-well cell-repellent surface microplates (Greiner BioOne) and sterilised with UV irradiation for 30 min at 254 nm. Quantities of different microparticle designs were determined to achieve a consistent matched surface area of 0.32 cm^2^/well for cell attachment across all 2D and 3D cultures. Following this, microparticles were conditioned in the relevant cell culture medium for 1 hour. Cells were then seeded at 6,000 cells/well across all experiments.

### 2.9 Morphological profiling of cell-microparticle aggregate morphology

MG-63 and U2OS cells were cultured on microparticles for 48 hours. Cell-microparticle aggregates were stained with Calcein AM (30002-T, Biotium) according to manufacturer instructions. Cell-microparticle aggregates were imaged using an EVOS M7000 microscope (ThermoFisher Scientific, United States), and a custom image analysis pipeline was developed and used for automated quantification of aggregate number and area across multiple fields of view using ImageJ (Figure 1).

**Figure 1.**
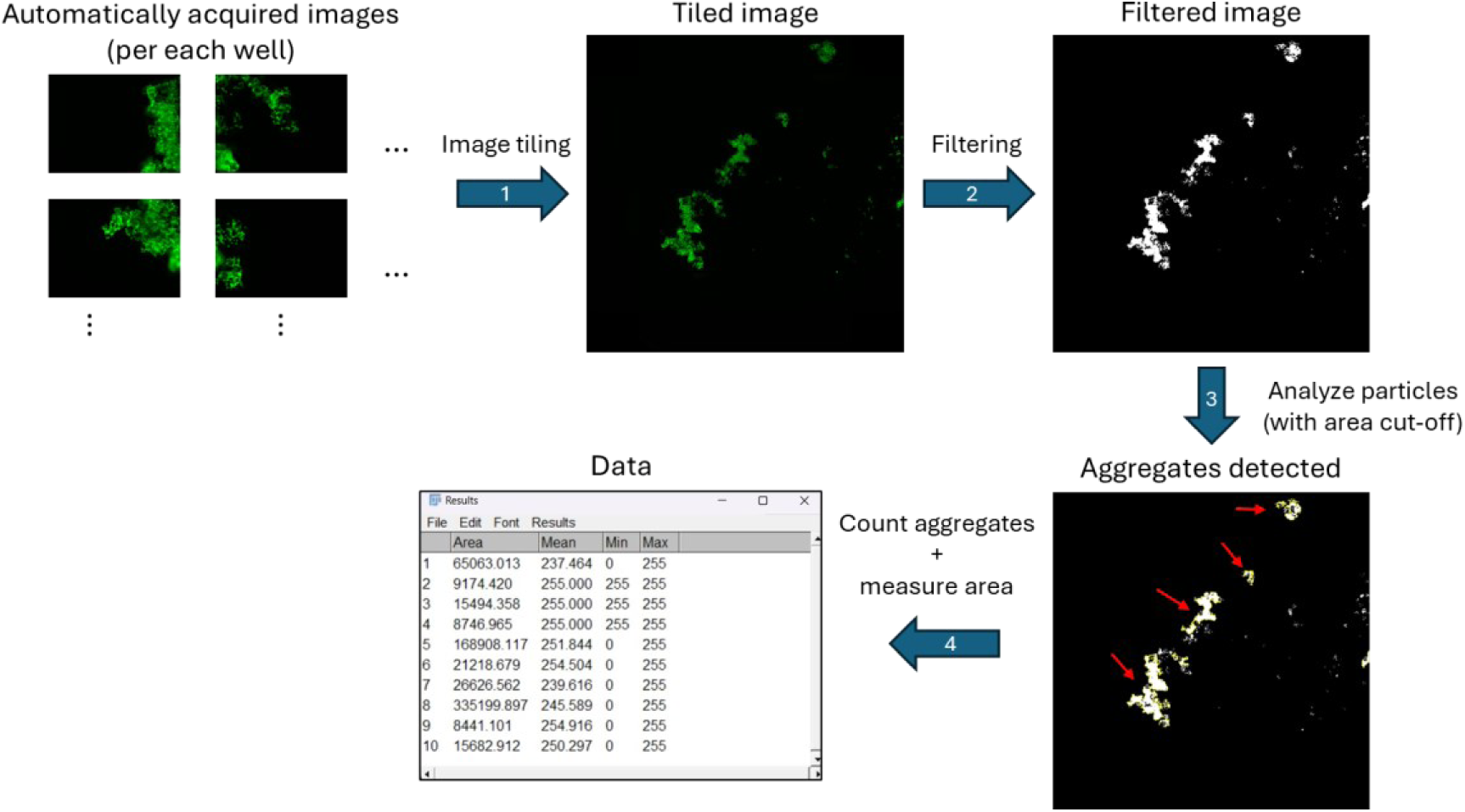
Schematic outlining cell-microparticle aggregate image analysis. (1) Fluorescence images were acquired on EVOS M7000 and tiled using the Invitrogen EVOS M7000 Imaging System software and analysis performed using ImageJ. (2) Otsu’s thresholding was applied, followed by median filtering (radius=10). (3) Aggregates were analysed with a minimal area threshold of 7584 µm^2^, corresponding to the maximum projected area of individual microparticles to ensure that only multicellular aggregates were included. Red arrows highlight the detected aggregates. (4) Aggregate numbers and sizes were measured using an ImageJ macro.

### 2.10 Doxorubicin treatment

MG-63 and U2OS cells were cultured for 24 hours, after which they were treated with 10 µM of doxorubicin (44583, Merck) for a further 72 hours. The intracellular distribution of doxorubicin could be visualized using fluorescence microscopy due to its inherent fluorescent properties.

### 2.11 Assessment of cell viability

MG-63 and U2OS cells were cultured on different microparticles for 96 hours. Cell viability was assessed using the ReadyProbes^TM^ Cell Viability Imaging kit (ThermoFisher Scientific, United States) and Calcein AM staining (30002-T, Biotium; PK-CA707-30002, PromoCell) to provide complementary readouts. Cells were imaged using the Zeiss LSM880 confocal microscope (Zeiss, Germany).

### 2.12 Assessment of metabolic activity

Following culture on microparticles for 96 hours, metabolic activity was assessed in untreated and doxorubicin-treated cell-microparticle aggregates, alongside their respective 2D controls, using the PrestoBlue™ Cell Viability reagent (ThermoFisher Scientific, UK). Briefly, culture medium was replaced with a 1:9 PrestoBlue™ to culture medium, then incubated at 37 °C for 1.5 hours for MG-63 cells and 1.75 hours for U2OS cells. Triplicate aliquots of the supernatant were assessed for fluorescence on a BioTek Synergy H1 microplate reader (Agilent, United States) at λ_exc_/λ_em_ of 560/590 nm.

### 2.13 Double-stranded DNA (dsDNA) quantification

Quantification of dsDNA of the cell lysates in untreated and doxorubicin-treated samples was measured using a Quant-iT™ PicoGreen™ dsDNA assay kit (Thermo Fisher Scientific, United States). DNA concentrations were calculated using a standard curve generated from the standards provided by the manufacturer. Fluorescence measurements were performed in doxorubicin-treated samples following extensive washing steps to remove residual doxorubicin.

### 2.14 Lactate dehydrogenase (LDH) cytotoxicity assay

Cytotoxicity induced by doxorubicin treatment was assessed by measuring LDH release into the culture medium. Following cell culture on microparticles for 96 hours, cytotoxicity was assessed in doxorubicin-treated cell-microparticle aggregates, as well as their untreated controls, using LDH-Glo™ Cytotoxicity Assay (Promega). LDH concentrations were calculated using a standard curve generated from the standards provided.

### 2.15 Untargeted metabolomics

MG-63 cells were cultured on microparticles as described above, and were either left untreated or were treated with 10 µM of doxorubicin (44583, Merck) after 24 hours of culture for a further 72 hours. At day 4 post-seeding, 150 µL of media per well were collected, centrifuged at 20,000xg for 5 minutes, and the supernatant was mixed with ice-cold acetonitrile (1:1) and vortexed briefly. After centrifugation, the supernatant was transferred to another tube and stored at 4 °C until used. Culture medium samples were processed in parallel as ‘media-only’ controls. Samples were then dried in a Speedvac centrifuge and reconstituted in 100 µl acetonitrile/water (5:1). Quality control (QC) samples were made by pooling 10 µl from each sample. Liquid chromatography-mass spectrometry analysis was performed using a SCIEX Exion LC system, coupled to a SCIEX 7600 ZenoTOF Q-TOF mass spectrometer with TurboV Optiflow ion source running a 50 µm ESI probe. The system was controlled by SCIEX OS (v3.0). A sample volume of 5 μL was injected onto a 50 μL sample loop with post-injection needle wash the same as the mobile phase start conditions. Injection cycle time was 1 min/sample. Separations were performed using an Agilent Poroshell 120 HILIC-Z column with dimensions of 150 mm length, 2.1 mm diameter and 2.7 μm particle size, equipped with a guard column of the same phase.

The mass spectrometer was operated in both positive and negative electrospray ionisation modes using the following source parameters unless otherwise specified: curtain gas pressure, 50 psi; source temperature, 400 °C; nebuliser gas pressure (GS1), 50 psi; heater gas pressure (GS2), 70 psi; and declustering potential, ±80 V. For positive ion mode (ion spray voltage 5500 V), mobile phase A consisted of water containing 10 mM ammonium formate and 0.1% formic acid, while mobile phase B was acetonitrile and water (9:1, v/v) with 10 mM ammonium formate and 0.1% formic acid. Separation was performed by gradient chromatography at a flow rate of 0.25 ml/min, starting at 98% B for 2 minutes, followed by a linear decrease to 5% B over 20 min, held at 5% B for 2 min, then back to 98% B. Re-equilibration time was 5 min. Total run time, including a 1 min injection cycle, was 30 min.

For negative ion mode (ion spray voltage, -4500 V), mobile phase A was water with 10 mM ammonium acetate adjusted to pH 9 with ammonium hydroxide and 20 µM medronic acid, mobile phase B was 85:15 acetonitrile and water with 10 mM ammonium acetate adjusted to pH 9 with ammonium hydroxide and 20 µM medronic acid. Separation was performed by gradient chromatography at a flow rate of 0.25 ml/min, starting at 96% B for 2 minutes, ramping to 65% B over 20 min, hold at 65 % B for 2 min, then back to 96% B. Re-equilibration time was 5 min. Total run time including 1 min injection cycle was 30 min.

Data was acquired in an information-dependent manner across 10 product ion scans, each with an accumulation time of 100 ms and a TOF survey scan with accumulation time of 250 ms. Zeno pulsing was used for both scan types. Total cycle time was 1.3 s. Isotopes within 4 Da were excluded from the scan.

Metabolite analysis was performed using Progenesis QI v. 3.0.7 (Waters Corporation, United States). Compound identification was done using HMDB, LipidMaps and NIST MS/MS libraries. Raw data was normalized to QC sample using Progenesis QI to correct for systematic analytical variation. Features with a relative standard deviation (RSD) greater than 30% across QC injections and interquartile range below 40% were excluded, and the filtered dataset was log_10_-transformed and auto-scaled. Multivariate analyses (PCA, PLS-DA, and oPLS-DA) were conducted in MetaboAnalyst 6.0. Variable importance in projection (VIP) scores, calculated from PLS-DA, quantify contributions of each variable to the observed separation between sample groups. VIP values greater than 1 were used as the cutoff values for quantitative enrichment analysis (QEA).

### 2.16 Statistics

Results are presented as mean ± standard deviation (SD). GraphPad Prism (version 9.3.1) was used for statistical analysis. The ROUT test was used to identify the presence of outliers (Q=0.5%), and data normality was assessed using the Shapiro-Wilk normality test (α=0.05) prior to further analysis using either non-parametric or parametric statistical testing, as appropriate. A *p*-value < 0.05 was considered statistically significant.

## 3. Results

### 3.1 DoE model predicts microparticle design parameters with high accuracy and informs design of surface topography

We first employed a Design of Experiments (DoE) approach to systematically generate and optimise topographically patterned microparticles with defined surface architectures. DoE approaches are widely used for the development and refinement of complex processes, with response surface designs being used to model quantitative relationships between experimental factors and measured outcomes. However, as the number of experimental factors increases, the feasibility of conducting a full factorial design diminishes rapidly due to the exponential growth in experimental runs required. To address this, it is often practical to prioritise main effects and low-order interactions. To our knowledge, the interactions between microparticle fabrication process parameters and their influence on topographically patterned microparticle architecture has not been elucidated.

To model the fabrication process (Fig. 2A), a Central Composite Design (CCD) of four factors and three levels was implemented, focusing on key fabrication parameters selected based on prior studies and known influence on emulsion stability and particle formation [37]: Combined concentration of PLA and FA, PLA concentration, homogenisation speed and stirring speed (Fig. 2B). The response outputs predicted were microparticle size, dimple size, and their respective PDI values (size distribution uniformity). Although the term ’topography’ broadly refers to surface features, we use the term ’dimpled’ throughout to describe our specific concave topographical designs for clarity and consistency. The model was trained on 52 runs, including duplicates. DoE analysis identified a significant dependency on homogenisation speed, concentration [PLA+FA] and concentration [PLA] as key determinants of microparticle architecture (Table 2). The combined effect of homogenization speed with total concentration of polymer and FA was also significant for microparticle design outcomes (Table 2). Among the factors tested, stirring speed did not significantly influence microparticle or dimple size nor PDI (*p*>0.05).

**Figure 2.**
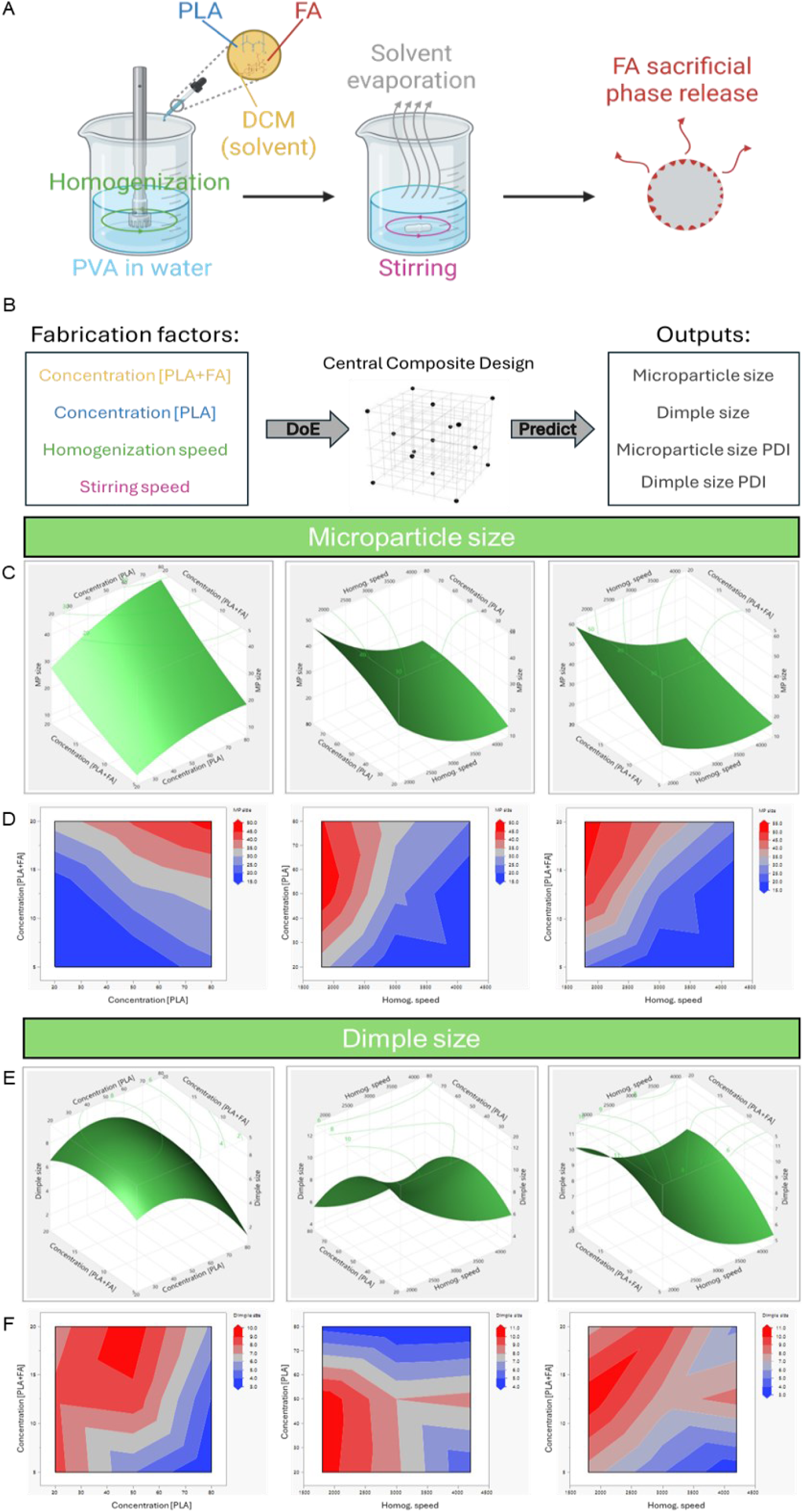
DoE model shows significant effect of fabrication parameters and their second-order interactions. (A) Schematic showing the microparticle fabrication process. (B) Schematic illustrating DoE model design. (C) Response surface plots showing effect of concentration [PLA+FA], concentration [PLA] and homogenization speed on microparticle size. Contour lines with predicted values are at the top plane of the plots. (D) Contour plots showing effect of concentration [PLA+FA], concentration [PLA] and homogenization speed on microparticle size. (E) Response surface plots showing effect of concentration [PLA+FA], concentration [PLA] and homogenization speed on dimple size. Contour lines with predicted values are at the top plane of the plots. (F) Contour plots showing effect of concentration [PLA+FA], concentration [PLA] and homogenization speed on dimple size. Abbreviations: PLA – polylactic acid, FA – fusidic acid, DCM – dichloromethane, PVA – poly(vinyl alcohol), DoE – design of experiments, PDI – polydispersity index, MP – microparticle.

**Table 2.** Effect summary report describing the top five significant fabrication factors and interactions for the entire DoE model. Effects are listed in order of significance (without information about their direction). Only sources with statistically significant effects were shown. LogWorth values above 2 corresponds to p<0.01. All p-values presented in this table are less than 0.00001.

| Source | LogWorth |
| --- | --- |
| Homogenization speed | 14.831 |
| Concentration [PLA+FA] | 14.752 |
| Concentration [PLA] | 12.263 |
| Homogenization speed*Concentration [PLA+FA] | 10.210 |
| Homogenization speed*Concentration [PLA] | 5.360 |

Contour and surface prediction plots demonstrate that topographical features could be tuned across a wide design space. Response surface plots and contour plots (Fig. 2C, D) revealed the influence of significant fabrication factors on microparticle size. Microparticle size can be reliably tuned by adjusting polymer content and homogenization speed, with diameter increasing steadily as either [PLA] or [PLA+FA] concentration rises, indicating that total PLA and FA content as well as the PLA fraction exert additive effects on microparticle size. On the other hand, dimple size is highly sensitive to interactions between formulation parameters, with non-linear and non-additive effects across all contour plots (Fig. 2E, F). Dimple formation is maximised at mid-range [PLA+FA] concentrations and lower homogenisation speeds, with intermediate values of [PLA] or total solids [PLA+FA] concentration often yielding the largest topography features. Dimple size decreased with increasing homogenisation speed overall (Fig. 2F). Response surface plots and contour plots also showed that higher levels of homogenization speed and lower levels of [PLA+FA] concentration increased microparticle size PDI, and lower PLA concentrations appear to compromise dimple uniformity by increasing the dimple size PDI (Fig. S1).

The robustness of this approach allowed us to generate microparticles across a broad morphological spectrum, from highly uniform spherical particles with minimal surface texture to particles characterised by one flattened side (Fig. 3A). The modelled design space enabled the rational selection of fabrication conditions to generate microtopographies with defined levels of uniformity. By systematically varying emulsion conditions, we produced topographically patterned microparticles with controlled diameter (ranging from 8-63 µm) and surface topographies (mean dimple size ranging from 2-13 µm) (Fig. 3A, B). The accuracy of the model in predicting design outputs was assessed, where actual values were plotted against predicted values. The DoE model showed high goodness of fit for all the outputs: microparticle size (R^2^=0.93), dimple size (R^2^=0.86), microparticle size PDI (R^2^=0.92) and dimple size PDI (R^2^=0.86) (Fig. S2). To validate the model, five additional validation runs were carried out (Fig. S3). Predictive ability was high for microparticle size (Q^2^=0.89), dimple size (Q^2^=0.73) and microparticle size PDI (Q^2^=0.78), indicating strong agreement between predicted and observed values. However, the model exhibited low predictive power for dimple size PDI (Q^2^=-18.69), indicating limited utility for predicting PDI under new conditions.

**Figure 3.**
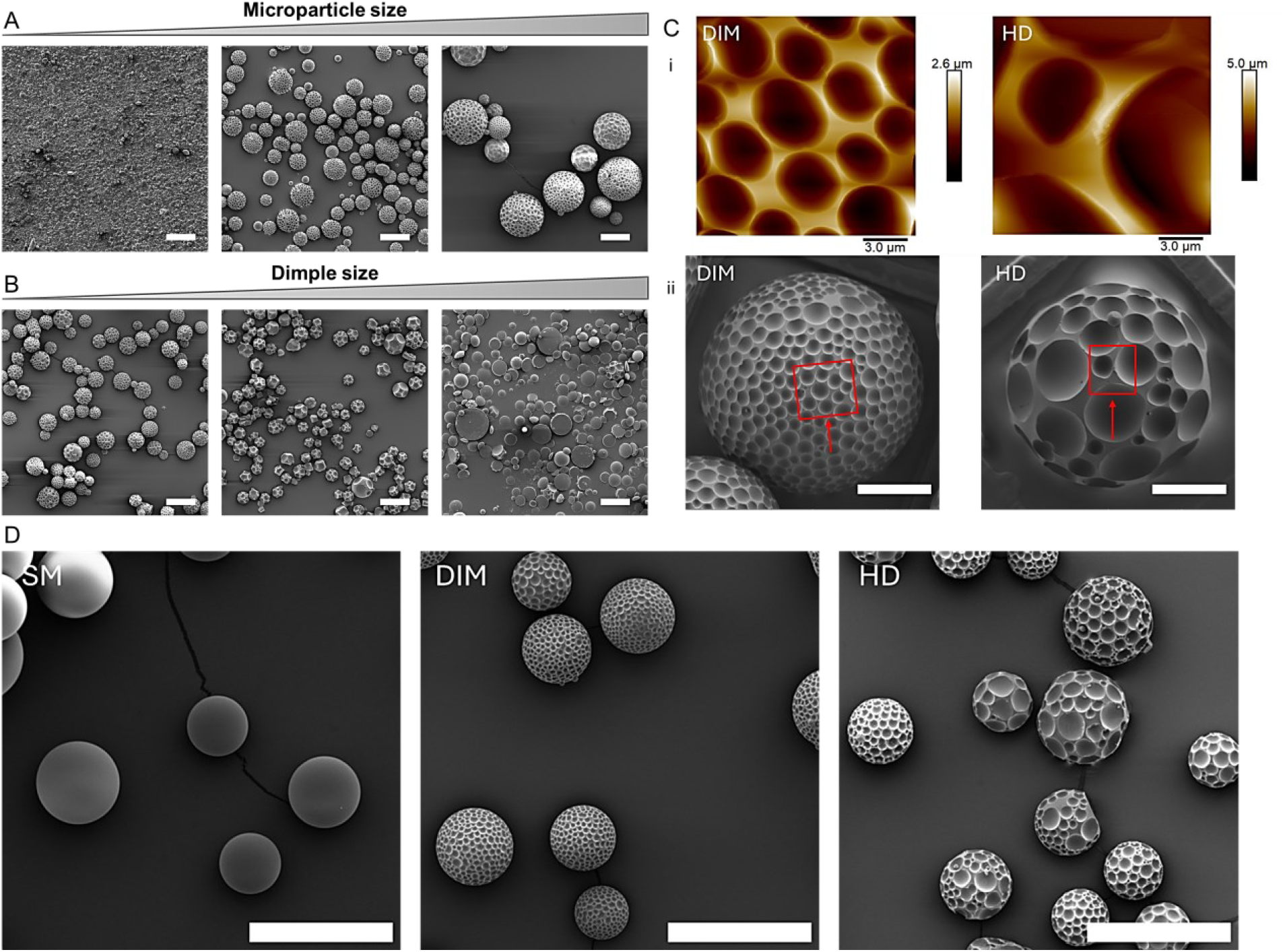
Range of topographically-engineered microparticles fabricated using the DoE model. Microparticles with varying (A) microparticle size and (B) dimple size. Scale bar: 50 µm. (C) Surface topographies of dimpled (DIM) microparticles, and heterogeneously dimpled (HD) microparticles visualised by (I) AFM and corresponding (II) SEM images of the same region of interest (highlighted by a red rectangle). Scale bar: 20 µm. (D) SEM images of microparticle designs selected for subsequent cell studies: smooth (SM), dimpled (DIM) and heterogeneously dimpled (HD). Scale bar: 100 µm. Abbreviations: SM – smooth, DIM – dimpled, HD – heterogeneously dimpled.

Using the developed RSM model, we next determined the optimal fabrication conditions for producing topographically textured microparticles with distinct size and dimple characteristics for cell studies. An optimization routine using the prediction profiler in JMP Pro 18 was applied to identify the optimal combination of factor levels that would yield microparticles meeting target criteria, namely particle diameter of ∼50 µm, in line with our previously published work on MSCs [35], with varying surface dimple sizes and PDI. To confirm the surface topography of different examples of the textured microparticles obtained using this model, SEM imaging and AFM measurements were conducted (Fig. 3C). Depth profiling revealed deeper dimples in the heterogeneously dimpled group, with a mean depth exceeding 2.63±0.7 µm relative to 1.86±0.4 µm in the uniform dimpled design (Fig. S4). It should be noted that AFM probe access to the full depth of some of the dimples was limited in the heterogeneously dimpled microparticles only, so the reported mean values may underestimate the true depth in this microparticle design. These measurements confirmed the presence of distinct topographical features, supporting our selection of microparticle design variants for downstream cell-material interaction studies. Dimple depth emerged *post hoc* from the selected fabrication parameters as distinct outcomes.

Together, this informed selection of a subset of particle conditions for downstream biological evaluation. Using the DoE model’s predictions, two topographically-textured microparticles (‘dimpled’ and ‘heterogeneously dimpled’) were fabricated, with ‘smooth’ microparticles as control (Fig. 3D). The uniformly dimpled particles displayed small, regular surface features (subsequently termed ‘dimpled’), and the more heterogeneously dimpled particles exhibited larger, less uniform dimple distribution (subsequently termed ‘heterogeneously dimpled’ to reflect increased topographical complexity). These were selected to investigate how microtopography and 3D spatial heterogeneity of topographical features influence osteosarcoma cell behaviour.

All three microparticle types exhibited comparable microparticle sizes and minimal surface microporosity (Table 3). The heterogeneously dimpled microparticles displayed increased dimple size with higher variability, as reflected by the higher dimple size PDI compared to the uniformly dimpled microparticles. While comparisons between dimpled and heterogeneously dimpled particles involve multiple varying features, which would not allow attribution of cell responses to a single parameter (i.e. dimple size versus depth), these categories enable an initial screen of how cells respond to distinct topographical surface environments, forming the basis for future studies to delineate individual topographical effects.

**Table 3.** Characteristics of microparticle designs used.

|  | **SM** | **DIM** | **HD** |
| --- | --- | --- | --- |
| **Microparticle size (µm)** | 59.07 ± 12.75 | 58.09 ± 15.47 | 58.39 ± 17.97 |
| **Dimple size (µm)** | N/A | 3.49 ± 0.96 | 6.52 ± 3.10 |
| **BET surface area (m<sup>2</sup>/g)** | 0.1393 | 0.1791 | 0.2183 |
| **V<sub>micro</sub> (cm<sup>3</sup>/g)** | 0.0000024 | 0.0000043 | 0.0000051 |
| **Microparticle size PDI** | 0.0466 | 0.0709 | 0.0947 |
| **Dimple size PDI** | N/A | 0.0757 | 0.2252 |

*Abbreviations: SM – smooth, DIM – dimpled, HD – heterogeneously dimpled*

### 3.2 Surface topography modulates osteosarcoma cell aggregation dynamics

To assess how microparticle surface topography influences osteosarcoma cell-microparticle interactions, aggregate size and number were measured for MG-63 and U2OS cells (representing fibroblastic and epithelial phenotypes, respectively) on the three microparticle types: smooth, dimpled, and heterogeneously dimpled microparticles. Aggregate formation reflects early cell-material interactions and their collective 3D responses, providing a functional readout of how surface features modulate cell aggregation, attachment, and spatial organisation.

MG-63 cells formed loosely organised cell-microparticle aggregates, with cells visible on the outer surfaces and inter-particle spaces of the aggregates (Fig. 4A). In contrast, U2OS cells appeared more deeply embedded between the microparticles, forming compact, enclosed aggregates, which is most visible in the smooth microparticle cultures (Fig. 4B). Microscopy images of MG-63 cells cultured on different microparticle designs (Fig. 4C) revealed visibly smaller aggregates on dimpled and heterogeneously dimpled particles compared to smooth microparticles. In contrast, U2OS cells showed no visible differences in aggregate morphology across microparticle types (Fig. 4D). Quantitative analysis confirmed that MG63 cells formed significantly smaller aggregates (Fig. 4E) than the no-substrate 3D controls, except for smooth microparticles (*p*=0.0839), as well as a significantly higher number of aggregates compared to the 3D controls (Fig. 4F; Fig. S5A). An increase in aggregate count typically reflects enhanced initial cell-microparticle adhesion and the nucleation of multiple separate aggregates. In contrast, changes in aggregate size tend to reflect subsequent cellular behaviours, such as proliferation, and may indicate differences in cell-cell interactions. While no statistically significant differences in aggregate size and count were observed between the different microparticle designs, the magnitude of deviation of the aggregate counts from the 3D no-material controls varied across the different topographical designs. The two topographically-textured microparticles displayed higher statistical difference from 3D controls than smooth microparticles, suggesting that surface topography may modulate aggregation behaviour to some degree.

**Figure 4.**
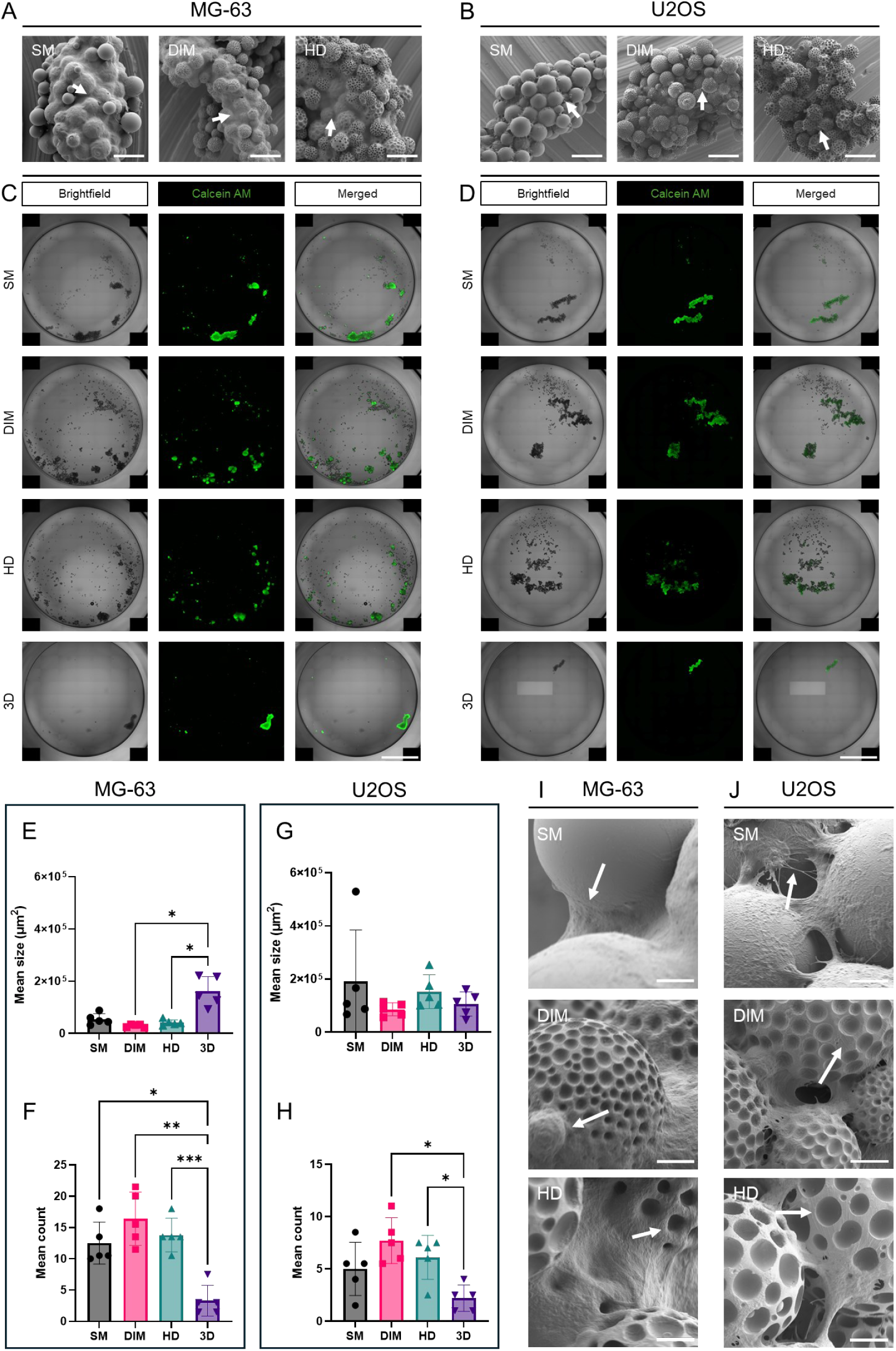
Cell-microparticle aggregate morphology depends on the cell phenotype and microparticle design. (A, B) SEM images showing MG-63 (A) and U2OS (B) cell-aggregate morphology at 96 hours post-seeding. Scale bar: 100 µm. (C, D) Tiled fluorescence microscopy image scan showing MG-63 cells (C) and U2OS cells (D) cultured on smooth (SM), dimpled (DIM) and heterogeneously dimpled (HD) microparticles at 48 hours. Scale bar: 2 mm. (E, F) Mean aggregate size (E; mean ± SD) and count (F; mean ± SD) of MG-63 cell-microparticle aggregates per well at 48 hours (n=5). (G, H) Mean aggregate size (G; mean ± SD) and count (H; mean ± SD) of U2OS cell-microparticle aggregates at 48 hours (n=5). (I) Representative SEM images of MG-63 cells cultured on SM, DIM and DD microparticles for 96 hours. Scale bar: 10 µm. Cells are indicated using white arrows. (J) Representative SEM images of U2OS cells cultured on SM, DIM and DD microparticles for 96 hours. Cytoplasmic protrusions are indicated using white arrows. Scale bar: 10 µm. RM one-way ANOVA with Tukey’s multiple comparisons test was performed. (* - p<0.05, ** - p<0.01, *** - p<0.001) Abbreviations: 3D – No-substrate 3D culture control, SM – smooth, DIM – dimpled, HD – heterogeneously dimpled.

Unlike MG-63 cells, U2OS cells showed no statistically significant differences in aggregate size relative to the no-substrate 3D controls and to one another (Fig. 4G, Fig. S5), which may indicate a more limited sensitivity to topographical modulation. However, they displayed a significantly higher number of cell-microparticle aggregates on topographically textured microparticles relative to the control, except for smooth microparticles (*p*=0.1232) (Fig. 4H). While the higher mean aggregate count was statistically significant for MG–63 cells cultured on smooth microparticles relative to the 3D control, the corresponding aggregate count increase in U2OS cells cultured on smooth microparticles did not reach statistical significance. Nonetheless, both cell lines showed comparable overall trends in aggregate counts (Fig. 4), implying a shared cellular response to 3D microparticles-based culture despite phenotypic differences.

To determine whether these early differences in aggregation translated to other changes in cellular response, we next examined cell morphology at 96 hours post-seeding. MG-63 cells displayed different morphologies on each of the three microparticle types (Fig. 4I), with cells appearing more spread out on smooth microparticles, and more rounded on the textured microparticles. On the heterogeneously dimples microparticles, cell membranes appeared to conform to the underlying dimpled surface features, extending into the dimples. U2OS cells, on the other hand, exhibited a spread out, flattened morphology across all microparticle designs. They also displayed long cytoplasmic protrusions across the microparticle surfaces, especially on the textured microparticles (Fig. 4J).

### 3.3 Cells cultured on 3D microparticles display reduced metabolic activity relative to 2D cultures

Quantification of cell numbers within 3D cell-microparticle aggregates using automated image analysis remains challenging due to the difficulty of reliable 3D nuclear segmentation within densely packed aggregates, as well as the limited imaging depth due to optical opacity of the PLA microparticles. To address this, assessment of metabolic activity (PrestoBlue) and total DNA content (PicoGreen) was performed within the same wells to investigate the effect of microparticle design on cell response. This dual-readout approach was used to distinguish between changes in cell number and cellular metabolism under varying culture conditions.

MG-63 cells cultured showed excellent viability across all microparticle designs (Fig. 5A) with significantly lower metabolic activity observed on all microparticle designs compared to 2D controls (Fig. 5B). However, dsDNA content was only significantly lower than the 2D controls in heterogeneously dimpled cultures (Fig. 5C). U2OS cells showed slightly reduced viability on all microparticle designs (Fig. 5D), with significantly reduced metabolic activity and cell number when cultured on microparticles relative to 2D cultures (Fig 5E, F). In contrast to MG-63 cells, U2OS cells cultured on smooth microparticles displayed a significant difference in metabolic activity to both topographically-textured microparticle designs, as well as to the conventional 2D cultures (Fig. 5E). Additionally, U2OS cells cultured on the heterogeneously dimpled microparticles exhibited more significant reduction in dsDNA content relative to the 2D cultures than those on smooth or dimpled designs (*p*<0.01 vs *p*<0.001; Fig. 5F).

**Figure 5.**
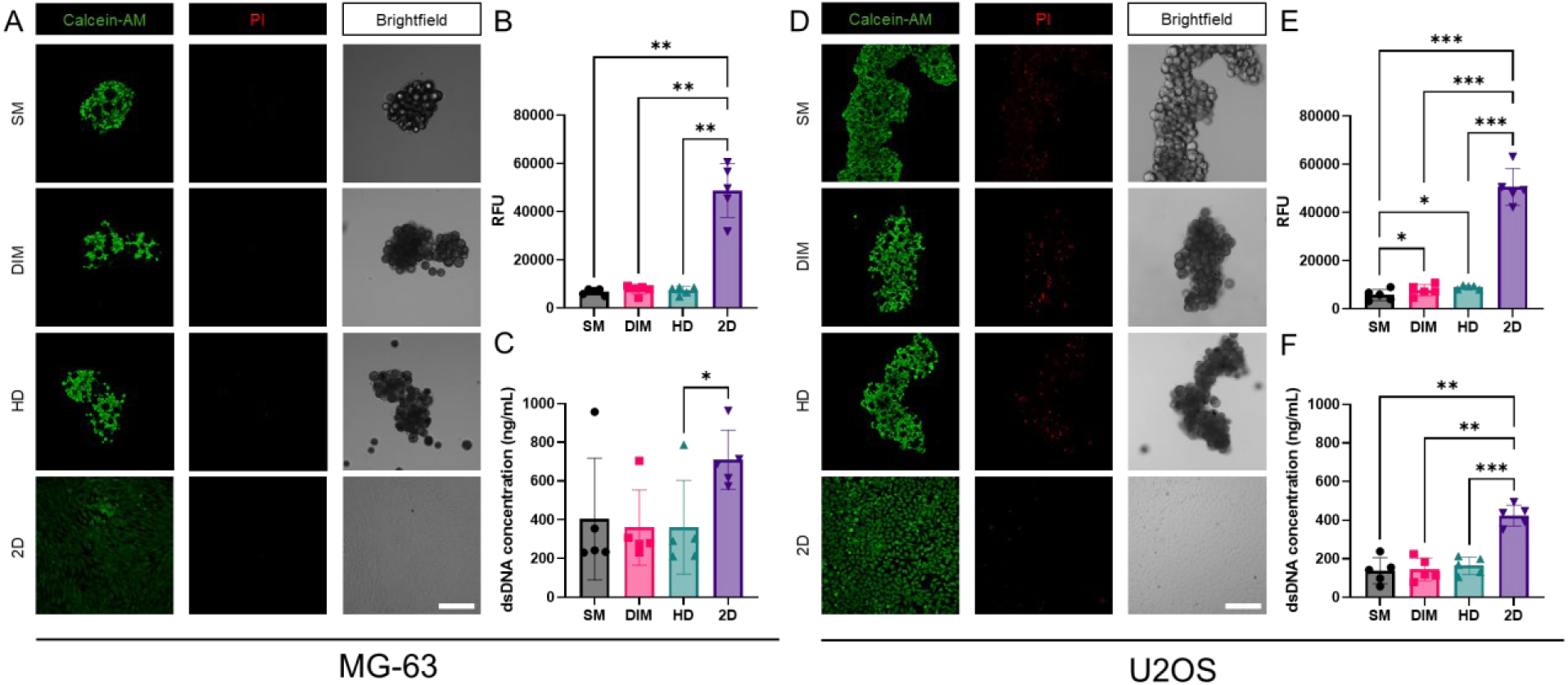
Cells cultured on microparticles exhibit significantly lower DNA content and metabolic activity relative to conventional 2D cultures, with differential responses between MG–63 and U2OS cells. (A) Representative maximum intensity projections of confocal microscopy images of MG-63 cells cultured on smooth, dimpled and heterogeneously dimpled microparticles for 96 hours (n=3), with a single optical plane shown for brightfield images for clarity. Scale bar: 2 mm. (B, C) Quantification of (B) metabolic activity using PrestoBlue (n=5, mean ± SD; RM one-way ANOVA with Geisser-Greenhouse correction and Tukey’s multiple comparisons test) and (C) dsDNA content by PicoGreen assay (n=5, mean ± SD; Friedman test with Dunn’s multiple comparisons test) performed on day 4 post-seeding. (D) Representative maximum intensity projections of confocal microscopy images of U2OS cells cultured on smooth, dimpled and heterogeneously dimpled microparticles at 6,000 cells/well for 96 hours (n=3), with a single optical plane displayed for brightfield images. (E, F) Quantification of (E) metabolic activity using PrestoBlue (n=5, mean ± SD, RM one-way ANOVA with Geisser-Greenhouse correction and Tukey’s multiple comparisons test) and (F) dsDNA content using PicoGreen (n=5, mean ± SD, RM one-way ANOVA with Geisser-Greenhouse correction and Tukey’s multiple comparisons test) assay at day 4 post-seeding. (* - p<0.05, ** - p<0.01, *** - p<0.001). Abbreviations: 2D – monolayer culture control, SM – smooth, DIM – dimpled, HD – heterogeneously dimpled.

Overall, U2OS cells cultured on microparticles exhibited higher statistically significant differences in DNA content and metabolic activity relative to 2D controls than MG–63 cells, indicating a greater sensitivity to microparticle-based culture, with additional phenotype-dependent modulation by surface topography. In both cell lines, the ratio of metabolic activity between 3D microparticle-based cultures and 2D controls was markedly lower than the corresponding ratio for dsDNA content (Fig. 5B, E). Given that no statistically significant differences in dsDNA content was observed across conditions on day 1 post-seeding (Fig. S6), indicating comparable initial seeding efficiency, this lower metabolic activity observed in 3D cultures may suggest a less proliferative or metabolically restrained state in 3D microparticle cultures compared to traditional 2D cultures.

### 3.4 Culture dimensionality and cell phenotype dominate response to doxorubicin treatment, with topography exerting a more subtle influence

To evaluate how osteosarcoma cells respond to chemotherapeutic stress in 3D microparticle cultures, cellular metabolic activity, total DNA content and cytotoxicity (using lactate dehydrogenase (LDH) release as a marker of membrane damage) were measured following exposure to 10 µM doxorubicin. Doxorubicin was chosen for its role as a first-line chemotherapeutic in osteosarcoma and its widespread use as a benchmark drug in cancer research, making it highly relevant for translational research. Measurements were taken on day 4 post-seeding, after 72 hours of drug exposure. Metabolic activity and dsDNA content were normalised to their respective untreated controls to enable comparison of relative drug sensitivity across culture conditions.

Doxorubicin treatment resulted in significant decreases in metabolic activity and dsDNA content in both MG-63 and U2OS cells across all culture conditions (Fig. S7). In MG-63 cultures, cell-microparticle aggregates remained structurally intact following treatment (Fig. 6A) and showed no significant increases in LDH content post exposure across all culture conditions (Fig. 6B), indicating limited cell membrane damage. When normalised to their respective untreated groups, MG-63 cells cultured on microparticles retained significantly higher normalised metabolic activity relative to their 2D counterparts (Fig. 6C). A significantly higher dsDNA content relative to 2D controls was only observed in dimpled microparticle cultures (Fig. 6D).

**Figure 6.**
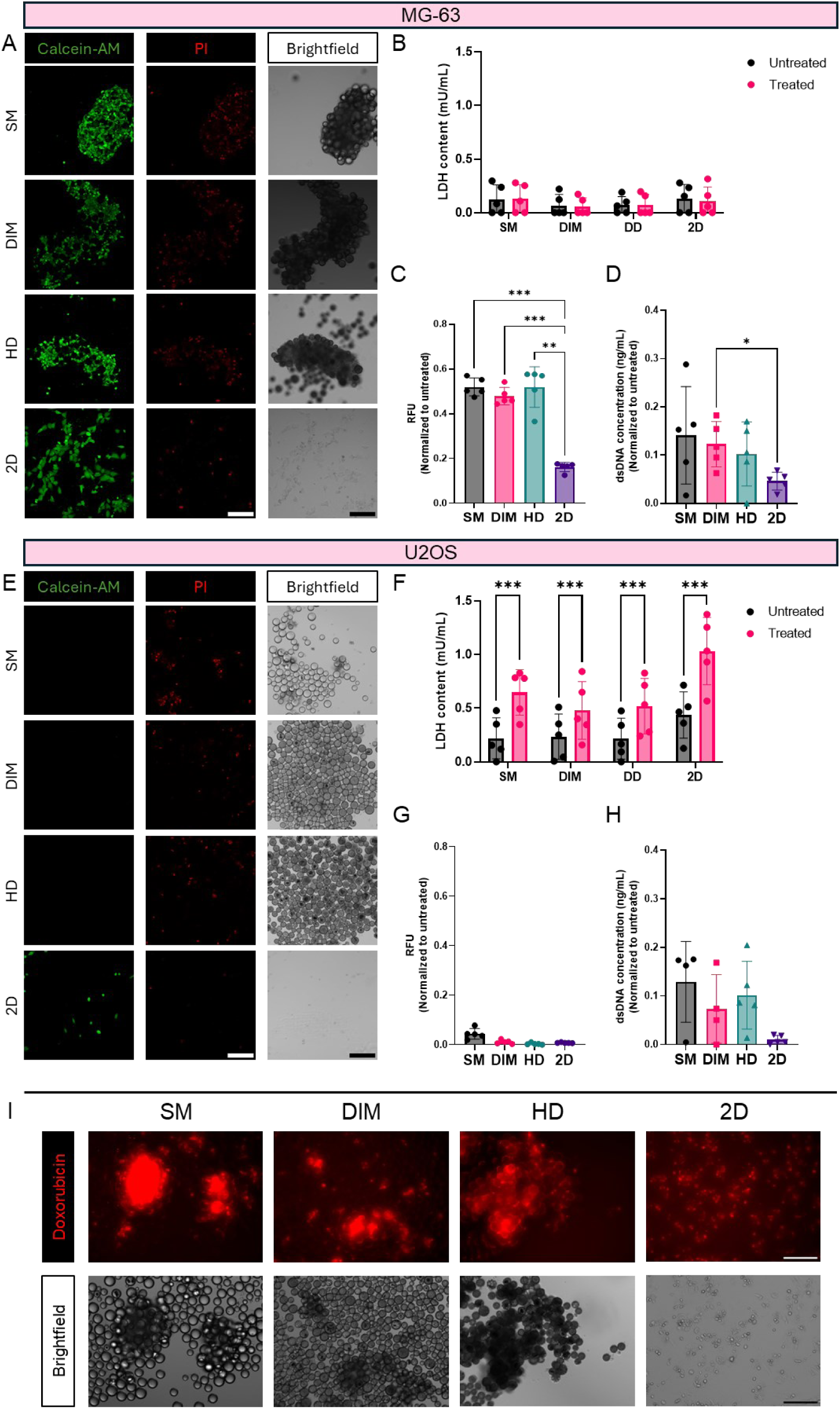
Cell response to doxorubicin treatment is primarily driven by cell phenotype and culture dimensionality, with lower influence of topography. (A) Representative maximum intensity projections of confocal microscopy images of MG-63 cells (n=3) cultured on smooth, dimpled and heterogeneously dimpled microparticles for 24 hours, followed by exposure to 10 µM doxorubicin for 72 hours. Brightfield images show a single focal plane corresponding to the confocal projection. Scale bar: 200 µm. (B-D) MG-63 cell response was measured using (B) LDH-Glo cytotoxicity assay (n=5, mean ± SD; two-way ANOVA with Tukey’s post hoc test), (C) PrestoBlue metabolic activity assay (n=5, mean ± SD; RM one-way ANOVA with Geisser-Greenhouse correction with Tukey’s multiple comparisons test) and (D) PicoGreen dsDNA quantification assay MG-63 (n=5, mean ± SD; RM one-way ANOVA with Geisser-Greenhouse correction with Tukey’s post hoc test). (E) Representative maximum intensity projections of confocal microscopy images of U2OS cells (n=3) cultured on smooth, dimple and heterogeneously dimpled microparticles for 24 hours then exposed to 10 µM of doxorubicin for 72 hours, with a single focal plane shown for brightfield images for clarity. Scale bar: 200 µm. (F-H) U2OS cell response was measured using (F) LDH-Glo cytotoxicity assay (n=5, mean ± SD; two-way ANOVA with Tukey’s post hoc test), (G) PrestoBlue metabolic activity assay (n=5, mean ± SD; RM one-way ANOVA with Geisser-Greenhouse correction with Tukey’s post hoc test) and (H) PicoGreen dsDNA quantification assay MG-63 (n≥4, mean ± SD; Kruskal-Wallis test with Dunn’s multiple comparisons test). (I) Representative maximum intensity projections of fluorescence imaging showing doxorubicin uptake by MG-63 cells under different culture conditions. Intracellular doxorubicin fluorescence was imaged on day 3 post-treatment. Scale bar: 200 µm. (* - p<0.05, ** - p<0.01, *** - p<0.001). Abbreviations: 2D – monolayer culture control, SM – smooth, DIM – dimpled, HD – heterogeneously dimpled.

In contrast, U2OS cells displayed heightened sensitivity to doxorubicin. U2OS cell-microparticle aggregates largely disaggregated following doxorubicin treatment, with widespread loss of cell-microparticle structures (Fig. 6E). LDH assays indicated significantly higher cytotoxicity in U2OS cultures across all conditions compared to MG-63 cells (Fig. 6F), consistent with the visible disassembly of U2OS aggregates. Metabolic activity was significantly diminished across all cultures when compared to their respective untreated controls (Fig. S7), showing no significant differences across all groups relative to 2D-cultured controls when normalised to their respective untreated controls (Fig. 6G). No significant difference was observed in normalized dsDNA content in U2OS cells (Fig. 6H).

We confirmed that PicoGreen fluorescence readouts remained linearly proportional to seeded cell number after doxorubicin treatment (R²>0.95 in both cell lines, n=3 each; data not shown), supporting the assay’s linearity and usability in doxorubicin-treated samples. To compare dsDNA content across conditions, treated values were normalised to the respective untreated controls. This approach assumes proportional DNA loss across cell densities; regression analyses confirmed this assumption in MG-63 cells (Q²=0.91; n=3) but showed greater variability in U2OS (Q²=0.61; n=3), confirming that the washing protocol mitigates interference, but results for U2OS should be interpretated with greater caution.

Deep doxorubicin penetration was observed within MG-63 cell-microparticle aggregates at day 3 post-treatment despite the structural stability of the aggregates (Fig. 6I). Together, these findings suggest that doxorubicin sensitivity is primarily influenced by cell line identity and dimensional culture context, with a smaller, context-specific influence of microparticle surface topographies.

### 3.5 Metabolomic profiling reveals culture dimensionality- and topography-driven changes in MG–63 cell metabolism

Untargeted metabolomic profiling was performed in MG-63 cultures to investigate topography-dependent metabolic adaptations, comparing smooth and dimpled microparticle cultures with and without doxorubicin treatment, to identify pathways associated with topography-mediated drug response (Fig. 7A). MG-63 cells, which have osteoblast-like properties, are known to respond to nano-etched membranes and patterned substrates [38], which makes them a useful model for assessing how microscale topography influences cellular metabolism. MG-63 cells also retained their 3D aggregate structure and a metabolically viable subpopulation across conditions, in contrast to U2OS cells. As U2OS 3D cultures lacked sufficient viability and structural integrity following doxorubicin treatment, they were not included in downstream metabolomic analysis.

**Figure 7.**
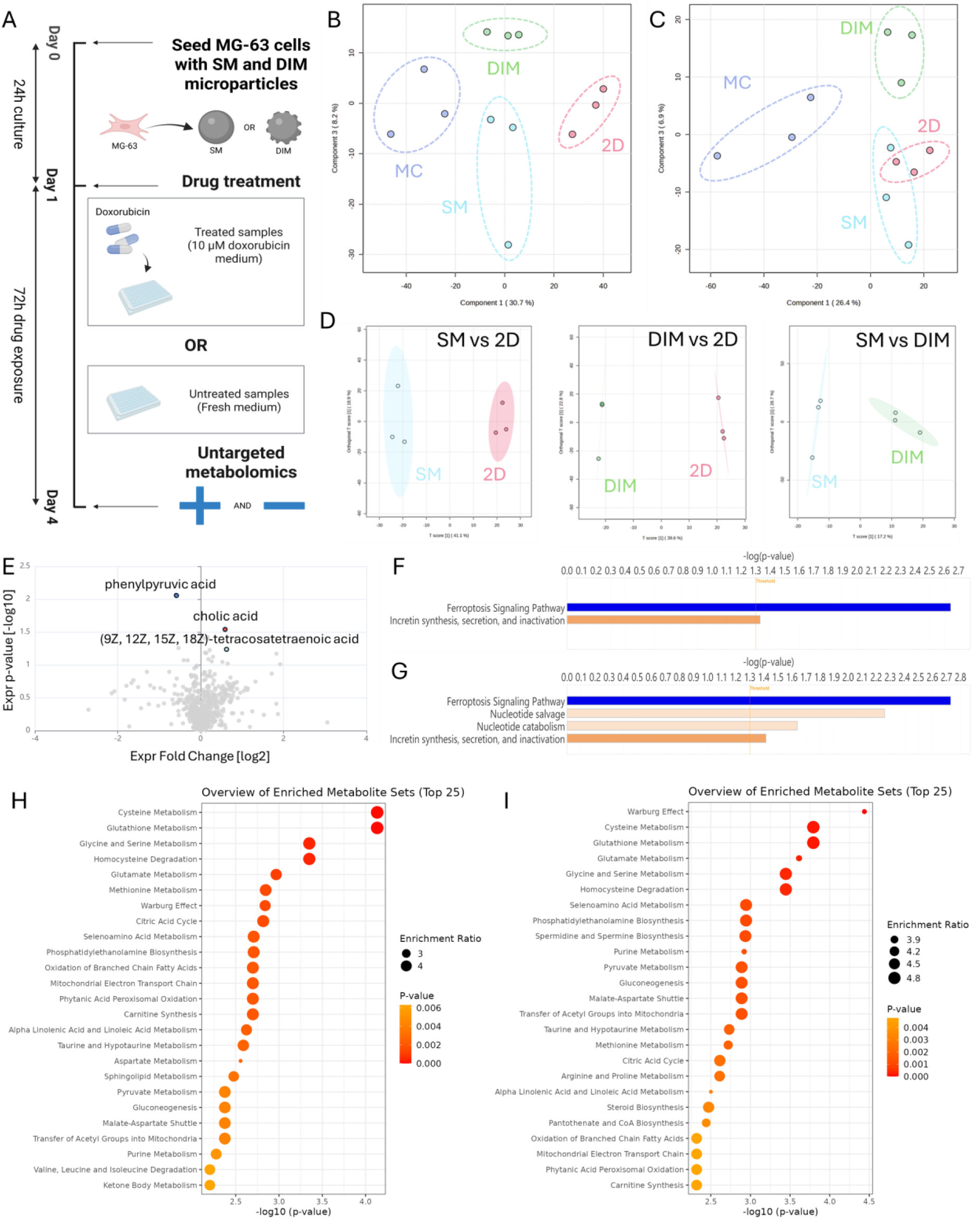
Metabolomic profiling reveals distinct metabolic signatures in 3D microparticles-based versus 2D cultures. (A) Schematic overview of experimental design: MG-63 cells were cultured on smooth and dimpled microparticles for 96 hours, with addition of 10 µM doxorubicin at 24 hours (n=3). (B, C) PLS-DA plots for the extracellular metabolic profiles in untreated (B) and doxorubicin-treated (C) cultures. (D) Orthogonal PLS-DA plots for the metabolic profiles in untreated samples comparing smooth versus 2D, dimpled versus 2D and smooth versus dimpled conditions. (E) Volcano plots showing the top differentially abundant extracellular metabolites in smooth versus dimpled microparticle cultures. (F, G) Ingenuity Pathway Analysis top canonical pathways for the differentially expressed metabolites in (F) smooth microparticle cultures versus 2D control cultures, and (G) dimpled microparticle cultures versus 2D. Positive z-score (orange) indicates predicted pathway activation, and a negative z-score (blue) represents predicted pathway inhibition. (H, I) Quantitative enrichment analysis performed for smooth microparticle versus 2D cultures (H) and dimpled microparticle versus 2D cultures (I). Multivariate and quantitative enrichment analyses were performed in MetaboAnalyst 6.0. Abbreviations: SM – smooth, DIM – dimpled, MC – media control

Partial least squares discriminant analysis (PLS-DA) revealed clear separation between untreated cells cultured on all microparticle designs, regardless of surface topography, and 2D controls (Fig. 7B), with less distinct separation between smooth microparticle cultures and 2D controls observed between doxorubicin-treated cultures (Fig. 7C). Smooth and dimpled microparticle cultures showed separation along Component 3 on the orthogonal PLS-DA (oPLS-DA) plots in both untreated (Fig. 7B) and treated (Fig. 7C) conditions, suggesting subtle topography-dependent metabolic differences. Unsupervised principal component analysis (PCA) revealed less separation between groups (Fig. S8), indicating that the observed differences in metabolic profiles are subtle and condition-dependent. Orthogonal PLS-DA plots of MG-63 cells cultured on smooth and dimpled microparticles showed clear separation from 2D control samples, indicating clearly altered metabolic states on microparticles relative to conventional 2D culture (Fig. 7D). While dimensionality of the culture (2D versus 3D) appeared to be the primary driver of metabolic divergence, a more subtle but reproducible distinction was also evident between smooth and dimpled microparticle cultures, supporting the potential role of 3D surface topography in modulating cell metabolism. Similarly, in the doxorubicin-treated samples, oPLS-DA analyses showed clear separation between 3D and 2D cultures, with a weaker separation between the two microparticle design cultures (Fig. S8C-E).

To investigate the metabolic pathways affected by surface topography, QEA was first performed to compare dimpled versus smooth microparticle cultures. No significantly enriched pathways were identified in this direct comparison, suggesting that the separation observed in the PLS-DA plot may reflect subtle, coordinated changes across multiple pathways rather than pronounced alterations in specific processes. This could also indicate involvement of low-abundance metabolites or fall below the resolution threshold of untargeted analysis. To explore topography-dependent changes at the individual metabolite level, metabolites with absolute fold change >1.5 and *p*<0.05 were investigated using Ingenuity Pathway Analysis (IPA), a curated bioinformatics tool for biological pathway inference. Only two metabolites were significantly altered meeting these thresholds: phenylpurvic acid was significantly reduced (FC=-1.51; *p*=8.64×10^-3^) and cholic acid significantly increased (FC=1.51; *p*=2.89×10^-2^), indicating that these two metabolites are particularly sensitive to topography in MG-63 osteosarcoma cells (Fig. 7E). Tetracosatetraenoic acid, a very long-chain polyunsaturated fatty acid (24:4) belonging to the omega-6 fatty acid family [39], was also substantially altered, although it did not reach statistical significance (*p*=0.052; Fig. 7E). These findings indicate that surface topography modulates discrete metabolic features rather than triggering broad pathway-level changes.

IPA was employed to identify the most relevant pathways that link the metabolic changes observed to biological processes. MG-63 cells cultured on smooth and dimpled microparticles both exhibited predicted negative activation of the ferroptosis signalling pathway (z-score=-1.00) and predicted positive activation of incretin metabolism (z-score=1.34) relative to 2D cultures (Fig. 7F, G), suggesting reduced engagement of oxidative stress-induced cell death pathways in 3D culture. In addition, a modest predicted positive activation of nucleotide salvage and nucleotide catabolism pathways was only observed in MG-63 dimpled microparticle cultures relative to 2D controls (z-score=0.45), indicating some engagement of nucleotide turnover or stress-adaptive recycling pathways unique to this topography (Fig. 7F).

Having explored differences between microparticle topographies, we next examined how each microparticle condition alters cellular metabolism relative to standard 2D culture. A total of 138 metabolites passed the cut-off values (*p*<0.05, |FC|>1.5) in both smooth microparticle versus 2D cultures (41 down and 97 up) and dimpled versus 2D culture comparisons (37 down and 101 up; Fig. S9). Analysis of upstream regulators, which is used to infer upstream molecules potentially responsible for the observed metabolic changes, identified choline kinase and epidermal growth factor in both smooth vs 2D cultures as well as dimpled vs 2D culture comparisons, suggesting shared regulatory mechanisms across 3D microparticle cultures. In contrast, 3-nitropropinoic acid was only affected in the smooth microparticle vs 2D culture (Table S1). Toxicological function analysis, which links toxicity endpoints and their potential causal associations with metabolites in the dataset, revealed predicted alkaline phosphatase activation in both smooth and dimpled microparticle culture comparisons with 2D controls (Tables S1 and S2).

Comparisons of the metabolic profiles of MG-63 cells cultured on both microparticle designs versus 2D cultures shared and topography-specific differences. QEA was subsequently performed based on metabolites with VIP scores >1.0, revealing distinct pathway-level differences where untreated smooth and dimpled microparticles were each compared independently to 2D cultures. While many enriched pathways overlapped between the two comparisons, consistently perturbed pathways identified in both comparisons included glycine and serine metabolism, cysteine metabolism and glutamate metabolism (Fig. 7H, I), suggesting shared responses to 3D microparticle-based cultures. However, each condition also exhibited distinct pathway enrichments within the top 25, such as sphingolipid and ketone body metabolism in smooth microparticle culture comparisons, and arginine/proline and steroid biosynthesis in dimpled culture comparisons against 2D cultures. To examine metabolite-specific changes, we compared individual compounds across conditions. In smooth microparticle vs 2D cultures, galactosylglycerol (*p*=0.036) and tetracosapentaenoic acid (*p*=0.036) were significantly lower compared to 2D cultures, whereas these were not significantly perturbed in dimpled microparticle vs 2D cultures. Conversely, higher differential abundance of D-galactose (*p*= 0.023) and acetoacetic acid (*p*=0.034), and lower levels of proline (*p*=0.0068) was observed in dimpled microparticle cultures relative to 2D cultures, none of which were significantly perturbed in the smooth microparticle comparison versus 2D culture. A subset of metabolites was significantly perturbed (*p*<0.05) in both comparisons, including L-cysteine, serine, taurine, 5′-methylthioadenosine, succinate and malic acid. These findings suggest that while dimensionality appears to be the dominant driver of metabolic change, distinct topographical features further modulate specific metabolite abundances and pathways.

Although nominally significant differences in metabolite levels were observed between the doxorubicin-treated groups, none remained significant after correction for multiple testing, with all false discovery rates (FDR) exceeding 0.05 (Fig. S8F, G).

## 4. Discussion

Native bone exhibits complex heterogeneous 3D topographies [40] that serve as a bioinstructive template. This study adopts a ’material-by-design’ approach to allow systematic optimisation of engineered microparticle properties for improved osteosarcoma modelling, facilitating precise control over particle characteristics for mimicking the TME. Our previous work demonstrated that topographically-textured PLA microparticles can induce mesenchymal stem cell osteogenesis without added biochemical factors via mechanotransduction [35, 36]. Building on this, we investigated whether similar microscale topographical cues influence osteosarcoma cell responses, offering insights into how surface architecture modulates cancer cell responses. In contrast to hydrogel-based osteosarcoma models, e.g. using alginate, these PLA microparticles more closely replicate the stiffness and 3D topographical features of the native bone microenvironment [35], providing a more physiologically relevant platform for preclinical modelling and therapeutic testing.

Among the four fabrication factors investigated, homogenization speed, total [PLA+FA] concentration, and concentration of PLA in the polymer mixture were the most influential (Table 2). Other fabrication parameters, such as buffer composition, may also influence microparticle morphology by modulating phase separation dynamics [41], potentially even post-evaporation in the solid state. Findings demonstrated that microparticle size can be tuned in a relatively stable and linear manner, with a concentration effect being evident for microparticle size outcomes. Lower PLA concentrations favoured the formation of smaller microparticles, likely due to the reduced viscosity allowing for more efficient mixing. On the other hand, dimple size appears to be more sensitive to the investigated fabrication parameters, and exhibit points of instability where minor deviations from the fabrication parameters can lead to markedly different outcomes. High shear and PLA levels appear to reduce dimple size, possibly by over-stabilising the emulsion interface. In addition, lower PLA concentrations compromise dimple uniformity as evidenced by lower dimple diameter PDI, likely due to increased phase separation due to insufficient polymer chain entanglement and reduced matrix cohesion during the process of dimple formation, both of which can impair surface feature resolution under dynamic emulsification conditions [42]. Overall, the contour plot patterns suggest non-linear interactions between the variables being assessed with regards to surface topography, indicating that simple linear relationships do not capture the full complexity of topographically textured microparticle formation. The DoE model demonstrated high goodness-of-fit for microparticle size (R^2^=0.93), dimple size (R^2^=0.86) and microparticle size PDI (R^2^=0.92), and displayed high predictive power on the validation set for microparticle size (Q^2^= 0.89), dimple size (Q^2^=0.73) and microparticle size PDI (Q^2^=0.78). However, while the R^2^ value for dimple size PDI was high (R^2^=0.86), the prediction power was notably low (Q^2^=-18.69), suggesting poor generalisability for this output. This may reflect the inherent stochasticity of dimple formation by phase separation under the selected parameters due to unpredictable microdomain formation behaviour with fusidic acid [37]. Small variations in local concentration or interfacial tension may disproportionately affect PDI, even if mean dimple size remains consistent. Moreover, differences in environmental conditions such as humidity or temperature may introduce non-linear effects not captured by the current model. While this limits the model’s utility for predicting the uniformity of dimples formed, the excellent predictability observed for dimple size and overall microparticle diameter demonstrates the utility of our DoE model and design framework. Further optimisation, including increased experimental replication or including more stringent control over environmental parameters, may improve model performance in terms of topographical uniformity in future studies. Recalibration and retraining are effective strategies to enhance model performance, as supported by studies across different fields [43]. Expanding the experimental design space to include additional process variables, such as solvent composition or surfactant type, may also improve model performance. Moreover, the use of more advanced approaches, e.g. machine learning algorithms capable of capturing non-linear interactions or mechanistic models that incorporate the underlying physics of dimple formation [37], can lead to more reliable and accurate predictions of dimple sizes and enhance predictive capability for this challenging parameter. While the successful prediction of microparticle and dimple size demonstrates the model’s utility, it also highlights the need for continued development in predicting secondary characteristics, such as dimple depth, that may be governed by different underlying mechanisms or require alternative modelling approaches.

Cell-microparticle aggregate formation may be governed by multiple biophysical cues, including microtopographical features that modulate cytoskeletal organization and integrin clustering [35, 36], and particle curvature that facilitates cell wrapping and 3D multicellular assembly [44]. Despite phenotypic differences between cell lines, MG-63 and U2OS cells exhibited similar trends in aggregate count. This may reflect how surface dimples influence early cell adhesion dynamics by altering spatial distribution of adhesion sites [45]. However, total available surface area was calculated to be equivalent across all microparticle designs based on BET surface area analysis, suggesting differences in aggregate formation cannot be explained by differences in contact area alone. The observed trend may instead reflect the modulation of integrin-based adhesion by dimpled topographies [35] or their influence on mechanosensitive signalling pathways, as reported previously for MSCs cultured on dimpled microparticles [36, 46]. Surface topography may also promote favourable ‘nucleation sites’ for initial cell contact, enabling stable aggregate formation before fusion into larger multicellular structures [47]. Cell aggregates have been shown to interact with microparticles during collective migration, with cells actively taking up 1 µm particles as they progress [48], suggesting that the physical presence and surface characteristics of microparticles can influence cellular behaviour beyond simple contact area considerations. Therefore, investigating whether the increased number of smooth microparticles, used at quantities calibrated via BET analysis to match the total surface area of dimpled microparticles, affects aggregate morphology could help elucidate the mechanistic basis of observed differences in cell-microparticle aggregate formation. The differential cell morphologies observed support this. Although integrin-mediated mechanotransduction pathways are impaired [49, 50], these differences in cell morphology may still influence early cell-material interactions. Spatial organisation of cells within aggregates represents a critical, but often underappreciated, variable in 3D in vitro models. MG-63 cells exhibited spreading on the surfaces of the microparticle assemblies, while U2OS cells were embedded more deeply within the interstitial spaces of microparticle aggregates. These differences in 3D cell architecture extends beyond simple positioning effects, as the mechanical microenvironment experienced by cells is shaped by their spatial context. For example, surface-positioned cells may experience different flow patterns and shear forces by media perfusion, whereas embedded cells may be more exposed to more curved, confined extracellular geometries, leading to shorter molecular diffusion distances and enhanced molecular sensing [51].

Both cell lines exhibited significantly lower metabolic activity in 3D microparticle cultures compared to 2D controls. The reduction in metabolic activity exceeded that of the corresponding lower dsDNA content, which aligns with previous observations of disproportionately reduced metabolic activity in tumour spheroids and 3D cultures relative to conventional 2D cultures. This is thought to partially recapitulate features of cancer cell quiescence [52]. This finding suggests that cells cultured on microparticles may adopt a metabolically quiescent phenotype despite maintaining proliferative capacity due to altered cell-matrix interactions or cytoskeletal tension [36, 53]. This may impact mechanotransductive signalling via the integrin-cytoskeleton axis, supporting the use of these topographically textured platforms to model dormancy-like phenotypes in vitro. Previous studies have shown that MG-63 cells exhibit lower proliferation rates when cultured as 3D spheroids compared to 2D monolayers [54]. Our findings extend these observations to topography-defined 3D cultures. Remarkably, U2OS cells cultured on smooth microparticles showed significantly lower metabolic activity on topographically-textured microparticles, which was not similarly observed in MG-63 cells. This cell line-specific response could be attributed to the less differentiated, more immature osteoblastic phenotype of U2OS cells [55], making them more susceptible to physical cues that favour quiescence. Additionally, our findings suggest that the impact of topographical design on osteosarcoma cells is cell line-specific, with statistically significant differences in metabolic activity observed between microparticle topographies in U2OS cultures but not in MG–63. This highlights the importance of engineered microarchitecture in shaping osteosarcoma cell response and suggests that microparticle-based platforms offer a customisable system for unravelling how matrix geometry influences cell-intrinsic responses. As initial seeding efficiency was comparable across conditions, the observed decrease in metabolic activity by day 4 likely reflects phenotypic adaptation to the 3D environment. While differences in metabolic activity may reflect a more quiescent phenotype in 3D culture, alternative explanations include altered mitochondrial function [56] or changes in cell cycle progression towards G_0_/G_1_ [57]. This customisable microparticle system offers a modular platform to study how biophysical stimuli contribute to cancer dormancy.

Our findings also suggest that dimensional culture context (i.e. 2D versus 3D) and cell phenotype play dominant roles in modulating drug response, with a relatively lower influence from engineered surface topography. Doxorubicin’s cytotoxicity is primarily attributed to its ability to induce DNA damage, leading to apoptosis and cell cycle arrest [58]. A combination of assays was employed to assess the impact of doxorubicin treatment on osteosarcoma cells on varying microparticle designs, providing a more comprehensive understanding of the influence of microparticle design on doxorubicin-induced cytotoxicity in osteosarcoma cells. Inherent differences in cell death kinetics were observed between the cell lines, especially in terms of LDH release, consistent with previous studies [59]. The differential response to doxorubicin may be attributed to differences in p53 status and associated apoptotic capabilities: U2OS cells possess functional p53 and can activate apoptosis pathways in response to DNA damage. In contrast, MG-63 cells harbour mutations that impair p53-dependent cytotoxicity [60]. This fundamental difference in DNA damage response machinery explains the contrasting drug sensitivities observed between the two cell lines. MG-63 cells in 3D cultures also showed significantly higher metabolic activity than 2D control post-treatment when normalised to their respective untreated controls. As normalized DNA content was largely similar across conditions post-treatment, this indicates that the remaining subpopulation of cells in microparticle groups is more metabolically active, which is often associated with chemoresistance [61]. Previous studies have reported that 3D osteosarcoma cultures show greater resistance to chemotherapeutic drugs compared to 2D monolayer cultures [62–64], partly due to the higher expression of calcium–activated potassium channels compared to 2D cultures [62]. In contrast, the significantly higher post-treatment normalised DNA content in U2OS cells on smooth microparticles relative to 2D cultures, which was not observed with topographically textured microparticles, indicate that topographical features may enhance drug sensitivity in U2OS cells, possibly through altered mechanotransduction pathways that influence p53-dependent apoptotic signalling. While doxorubicin intercalates into DNA and may theoretically compete with PicoGreen for binding [65], the assay was performed on samples after extensive washing steps to remove unbound doxorubicin. While the residual interference from irreversibly bound doxorubicin cannot be fully ruled out, it is unlikely to significantly affect DNA quantification under the conditions used. Moreover, assay reliability was supported by post-treatment calibration experiments, where PicoGreen fluorescence remained linearly proportional to seeded cell number (R² > 0.95; MG–63 Q² =0.91, U2OS Q²=0.61). The results supported the validity of the DNA readout, while also highlighting the need for cautious interpretation in U2OS cultures. The lower Q² in U2OS may reflect the intrinsic heterogeneity in this line, consistent with previous reports of phenotypic variability, even among clonal derivatives [66].

The distinct clustering of metabolic profiles between 2D and 3D culture conditions indicates that dimensionality exerts a dominant influence on MG-63 cell metabolism, while surface topography contributes more subtly. No significantly enriched pathways were identified in the direct comparison between dimpled and smooth microparticles, suggesting that topography may introduce subtle changes across multiple pathways rather than pronounced alterations in specific metabolic processes and/or involve low-abundance metabolites. Analysis of individual metabolite profiles revealed highly specific topography-associated differences. Cholic acid and phenylpyruvic acid were the only significantly altered metabolites between the two microparticle designs, along with a notable change in tetracosatetraenoic acid just below significance threshold, reflecting lipid and phenolic metabolism driven by topographical features. Differences in phenylpyruvic acid levels in MG-63 cells on dimpled substrates suggests altered phenylalanine metabolism [67]. Elevated phenylpyruvate has been linked to impaired osteogenic differentiation of MSCs [68] and pathological states such as phenylketonuria, and has been noted as a metabolic risk factor in aggressive cancers [69, 70].

Therefore, its decline here may indicate a shift towards a more anabolic state, where MG-63 cells on dimpled surfaces might assimilate phenylalanine more efficiently. The diminished extracellular phenylpyruvate in the dimpled cultures may therefore signify a shift towards a more differentiated metabolic phenotype. This aligns with its osteoinductive properties seen with human MSCs [35, 36]. The accumulation of cholic acid extracellularly suggests that MG-63 cells on dimpled substrates may be secreting more cholic acid, indicating altered metabolite transport across the cell membrane. MG-63 cells have been reported to take up exogenous bile acids, e.g. deoxycholate, from serum-containing medium [71]. Topography-driven activation of Hedgehog signalling may be linked to this change and is closely linked to cholesterol metabolism [72], with PTCH1 receptor directly contributing to cholesterol efflux [73] leading to increased export of cholesterol-derived metabolites such as cholic acid. These findings may suggest a metabolic shift toward higher cholesterol processing, linked to the reported activation of Hedgehog signalling in MSCs on dimpled microparticles [36, 74]. Moreover, cancer cells commonly reprogram fatty acid metabolism, and very long-chain fatty acids have been shown to be elevated in cancer [75, 76]. Fatty acid metabolism plays a critical role in osteosarcoma development, and a prognostic risk score system has been developed based on fatty acid metabolism-related genes [77]. This is consistent with the role of ELOVL1 (elongation of very long-chain fatty acids protein 1) in cancer [75], responsible for synthesizing very long-chain fatty acids, and this fatty acid pool affects ferroptosis sensitivity [78]. Unlike broader metabolic perturbations expected from generalised cellular stress, the isolated significant changes in these three metabolites indicates targeted response pathways that are particularly sensitive to topographical microenvironmental cues and supports that physical cues can fine-tune metabolic pathways. These metabolites may serve as biomarkers for mechanotransduction-induced metabolic reprogramming in osteosarcoma cells, though the statistical significance of tetracosatetraenoic acid in dimpled versus smooth microparticle cultures (*p*=0.052) did not reach conventional significance threshold.

By focusing on total extracellular metabolite levels under defined conditions, our approach prioritises comparability between system-level outputs across different culture conditions. This enabled the identification of topography- or dimensionality-dependent trends in overall metabolic engagement, which are critical parameters when assessing the suitability of 3D culture systems for 3D cancer modelling or drug response studies. Traditional normalisation strategies (e.g. per DNA content) may misrepresent functional capacity in 3D systems due to differences in cellular engagement with the matrix and inherent spatial heterogeneity, where nutrient and oxygen gradients may lead to distinct subpopulations compared with the relatively uniform metabolic activity of monolayer cultures [79]. This complicates direct comparison of metabolite abundance across dimensional platforms, as equal cell numbers may not reflect equivalent metabolic output. In the context of tumour model development, absolute metabolic output more closely reflects clinical reality, where bulk tissue metabolic profiles, not per-cell activity, define therapeutic response. In addition, PrestoBlue^TM^ readouts were found to significantly vary between 2D and 3D conditions, reflecting differences in metabolic activity influenced by culture conditions. Moreover, accurate cell or nuclear segmentation in 3D cultures remains technically challenging, further limiting the reliability of normalised data. Therefore, total metabolic output was considered a more appropriate and translatable readout, preserving the integrity of the engineered system and capturing aggregate-level metabolic function, including the contributions of viable, proliferating and quiescent subpopulations. This also highlights the need for standardized approaches to enable cross-platform metabolomic comparisons. While real-time metabolic flux measurements (e.g. using Seahorse XF technology) could offer additional insights into each culture system’s metabolic landscape, these approaches are not readily compatible with microparticle-based 3D cultures [80] and were therefore not pursued in this study.

Consistent with emerging evidence that biomaterial cues can reprogram metabolism [81], we found a shared set of perturbed pathways present in both smooth and dimpled microparticle cultures when compared against 2D monolayer cultures. In both comparisons, L-cysteine, serine and taurine levels were elevated, while malic acid, succinate and 5′-methylthioadenosine (MTA) levels were significantly reduced. L-cysteine and serine amino acids support the synthesis of glutathione [82, 83], a major antioxidant and a key player in maintenance of redox homeostasis [84]. Taurine is an end-product of cysteine catabolism [85], and elevated levels indicate cellular adaptation to oxidative stress [86]. The reduction in MTA, a precursor in methionine salvage [87], may reflect increased methionine dependency in 3D cultures. Methionine addiction refers to the phenomenon in which cancer cells exhibit an increased requirement for methionine to sustain elevated transmethylation activity [88], and methionine deficiency in osteosarcoma cells has been shown to inhibit cell growth and induce impairment of mitochondrial functions [89]. Elevated succinate levels in 2D cultures, an oncometabolite shown to be tumorigenic in high levels in the TME [90], may reflect a more tumorigenic phenotype. Although malate itself has not been directly implicated in osteosarcoma, malic enzymes (MEs) catalyse its conversion to pyruvate while generating NADPH, crucial for maintaining cellular redox homeostasis and supporting the synthesis of macromolecules such as lipids [91]. Given that MG-63 cells are p53-deficient, and p53 regulation of MEs [92], the lower malate levels observed in the microparticle groups may reflect increased MEs activity, suggesting enhanced reductive biosynthesis in 3D cultures. Ingenuity Pathway Analysis revealed predicted downregulation of ferroptosis-related pathways, a form of programmed cell death characterized by iron-dependent lipid peroxidation, in both microparticle cultures. Ferroptosis has been linked to the regulation of invasion of osteosarcoma cell lines [93], which may suggest that microparticle-based 3D cultures confer protection from lipid peroxidation-induced cell death. Incretin-related pathways were also upregulated in both comparisons. While the role of incretins in osteosarcoma is not well understood, glucagon-like peptide-1 (GLP-1) has been reported to enhance the MG-63 cell viability [94] and promote synthesis of c-Fos expression, a proto-oncogene and a bone development regulator [95]. One notable difference was the predicted moderately positive pathway activation of nucleotide metabolism and salvage in the dimpled microparticle cultures versus 2D controls, which was not observed in the smooth microparticle comparison. This suggests that dimpled topography may slightly stimulate the rate of nucleotide turnover in MG-63 cells, potentially reflecting increased DNA repair activity and recycling of nucleotides [96]. When examining upstream regulators, IPA predicted the activation of choline kinase alpha (ChoK) in both microparticle cultures relative to 2D controls, which has been associated with drug resistance and metastasis [97]. Only the smooth microparticle vs 2D comparison showed potential activation of 3-nitropropionic acid (3-NPA), a mitochondrial toxin that reduces ATP synthesis and inhibits succinate dehydrogenase [98], suggesting impaired mitochondrial activity in smooth microparticle cultures. The differential abundance of galactosylglycerol and tetracosapentaenoic acid in smooth vs 2D cultures may reflect changes in membrane lipid composition or fatty acid turnover associated with 3D culture. By contrast, dimpled microparticles induced significant changes in D-galactose, proline, and acetoacetic acid, suggesting substantially altered carbohydrate and amino acid metabolism which is not observed to the same degree in smooth microparticle cultures. These observations support our conclusion that while the transition from 2D to 3D culture exerts the largest metabolic effect, microscale surface topography can be used to further tune specific metabolic pathways.

Together, these findings highlight that topography-induced metabolic differences were subtle, while culture dimensionality (i.e. switching from 2D to 3D) triggered much larger metabolic reprogramming, with broad pathway alterations. This is consistent with previous reports of over 1700 differential metabolites between 2D and 3D OS cultures, including ferroptosis [99]. The subtle differences observed between smooth and dimpled conditions in the context of 2D comparisons may underscore the need for future studies with targeted metabolomics or time-resolved analysis to better resolve the specific pathways engaged by surface topography. Yang *et al* have also reported that culture method makes a significant difference on metabolite levels in osteosarcoma cells, and similarly, ferroptosis signalling pathway was implicated in 3D-cultured SaOS-2 cells [99]. Moreover, Wang *et al* have demonstrated that serine and glycine metabolism promotes MG-63 cell proliferation in 2D cultures [100], suggesting that microparticles provide a permissive environment for anabolic amino acid metabolism, important for osteosarcoma proliferation.

The purpose of this study was to evaluate how 3D microenvironmental topography shapes osteosarcoma cell behaviour, drug response, and metabolic state. While dimensionality was the primary determinant of osteosarcoma responses, topographical features provided a secondary, cell line-dependent layer of modulation. The most consistent effects of topography were observed in aggregation dynamics, with subtler influences on proliferation, cell morphology and chemosensitivity. Rather than driving broad metabolic reprogramming, topography elicited metabolite-specific changes, with significant differential expression of phenylpyruvic acid and cholic acid in MG-63 cells. Together, these findings indicate that while dimensionality fundamentally alters osteosarcoma cell behavior, topographical variation refines our ability to dissect osteosarcoma heterogeneity and cell-matrix interactions.

## 5. Conclusion

This material-by-design workflow provides a robust and tuneable 3D platform for modelling the osteosarcoma TME. By enabling precise control over 3D surface topography and dimensionality, these engineered substrates recapitulate the stiffness of bone and address key limitations of traditional spheroid and hydrogel-based models. They provide a physiologically relevant and scalable in vitro system that captures key hallmarks of osteosarcoma, including the tendency of tumour cells to form clusters and heterogeneous drug responses. Our platform addresses a fundamental gap in osteosarcoma research by enabling systematic investigation of how ECM-inspired physical cues influence therapeutic resistance. This platform therefore offers a valuable tool for improving the predictive power of preclinical drug screening while reducing reliance on animal-derived materials.

## Acknowledgements

This study was funded by the Academy of Medical Sciences Springboard Scheme [SBF008\1057]. This work was supported by the Henry Royce Institute for Advanced Materials, funded through EPSRC grants EP/R00661X/1, EP/S019367/1, EP/P025021/1 and EP/P025498/1. We thank Shahla Kahn from the Analytical Services and Instrumentation Lab. The Bioimaging Facility microscopes used in this study were purchased with grants from BBSRC, Wellcome, Walgreen Boots alliance and the University of Manchester Strategic Fund. We also thank Steven Marsden and James Bagnall for their help with microscopy. Figures 2A and 7A were created in BioRender.

## Data Availability Statement

Data supporting the findings of this study will be made available upon reasonable request.

## Notes

### Competing Interest Statement

The authors have declared no competing interest.

