## Supplementary data for "A Design-of-Experiments Strategy for Engineering 3D Topographical Features in Osteosarcoma Modelling"

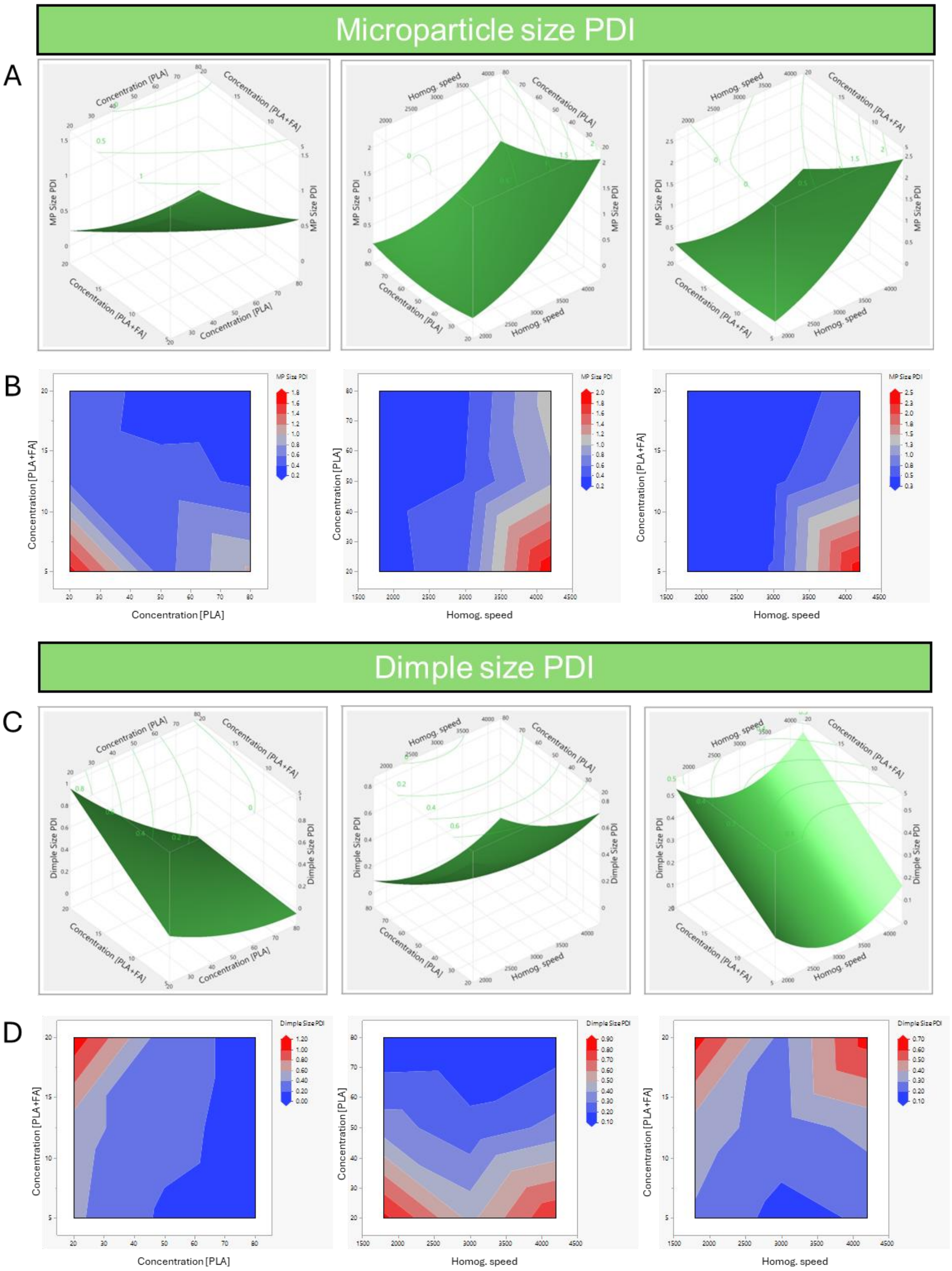


***Figure S1. Prediction plots for microparticle size polydispersity index (PDI) and dimple size PDI as a function of three fabrication factors: homogenization speed, [PLA+FA] concentration and [PLA] concentration.*** *(A) Surface plots and (B) contour plots for microparticle size PDI. (C) Surface plots and (D) contour plots for dimple size PDI.*
*Abbreviations: MP – microparticle, PDI – polydispersity index.*

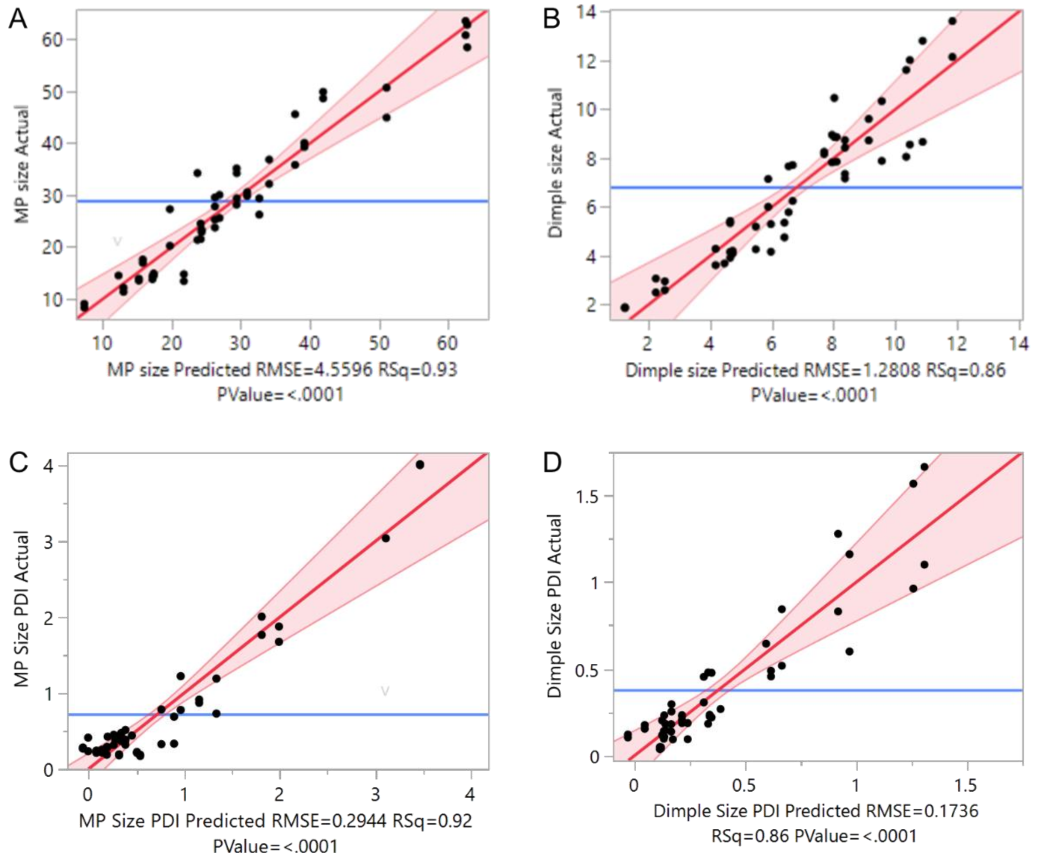


***Figure S2. Internal validation of the DoE model.*** *Prediction plots showing actual and predicted (A) microparticle size, (B) dimple size, (C) microparticle size PDI and (D) dimple size PDI.*
*Abbreviations: MP – microparticle, PDI – polydispersity index.*

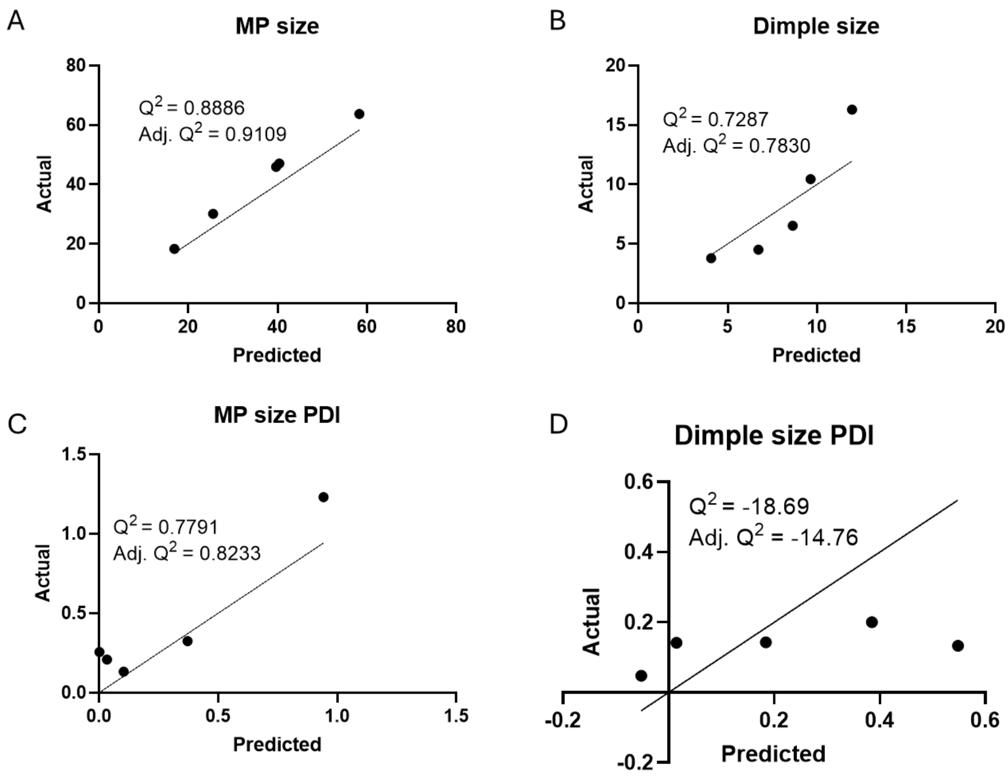


***Figure S3. Validation of the DoE model based on microparticle size and dimple size.*** *Validation of (A) microparticle size, (B) dimple size, (C) microparticle size PDI and (D) dimple size PDI. Black line represents the line of identity, with $Q^2$ calculated using the validation set mean.*
*Abbreviations: MP – microparticle, PDI – polydispersity index.*

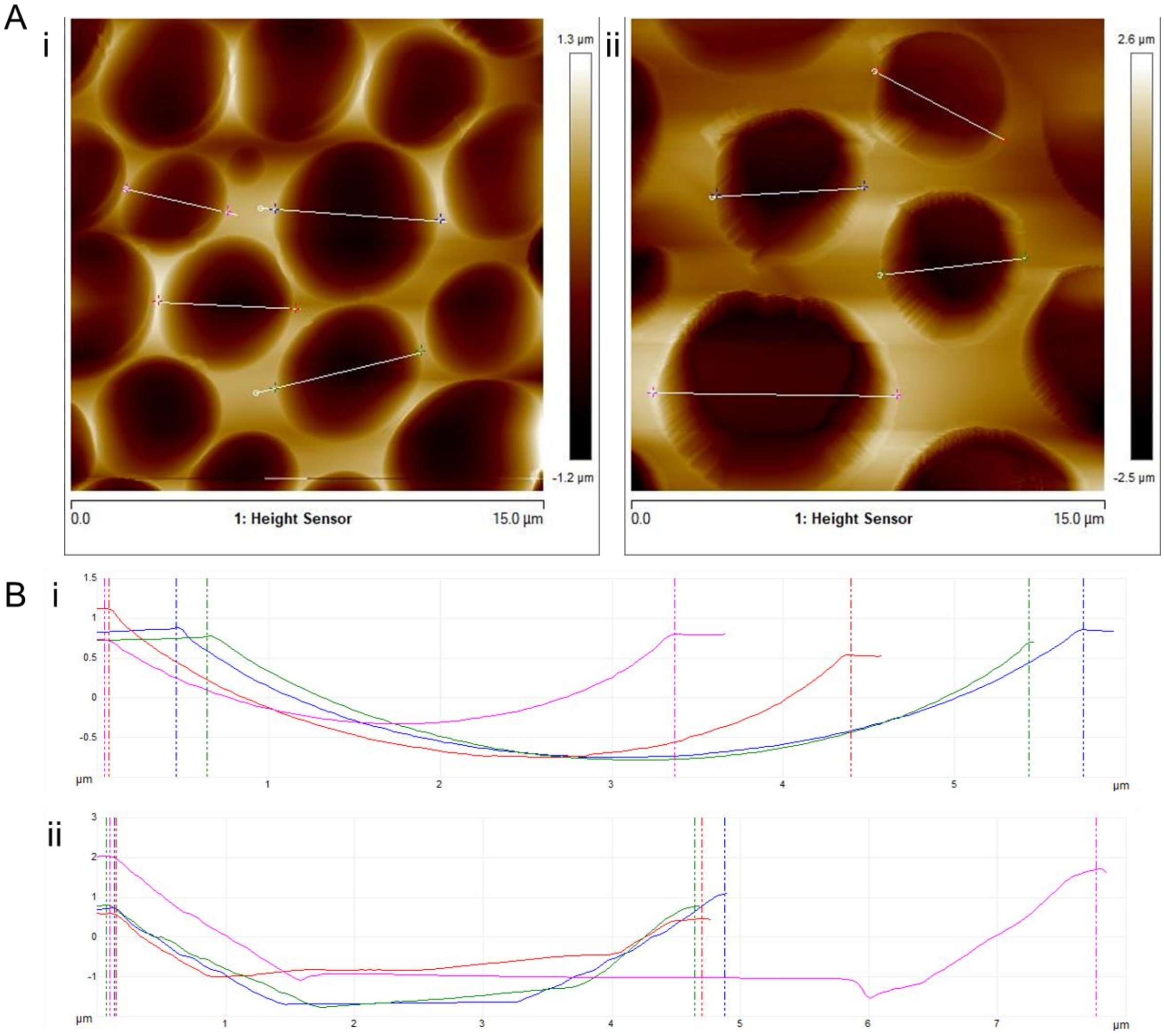


***Figure S4. AFM topography and depth profiles of dimpled and heterogeneously dimpled microparticles.*** *(A) Representative AFM images of (i) dimpled and (ii) heterogeneously dimpled microparticles. (B) Dimple depth profiles generated using Nanoscope Analysis (v3.0) software (n=16 dimples in ≥4 individual microparticles), showing profiles from representative (i) dimpled and (ii) heterogeneously dimpled microparticles.*

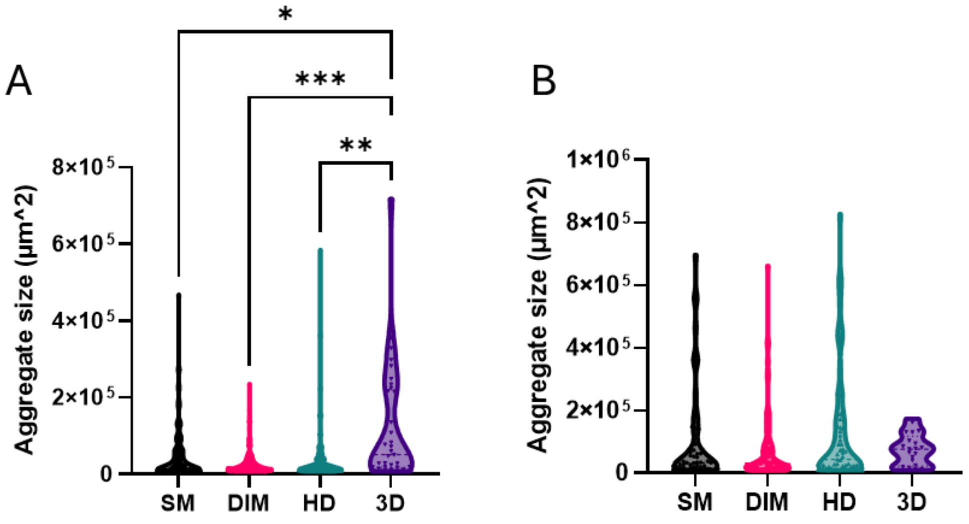


***Figure S5. Violin plots showing the size distribution of individual cell-microparticle aggregates in MG-63 and U2OS cells.*** *Size of cell-microparticle aggregates in (A) MG-63 cells and (B) U2OS cells. Each point represents one aggregate. Kruskal-Wallis test with Dunn's multiple comparisons was performed (* - p<0.05, ** - p<0.01, *** - p<0.001)*
*Abbreviations: 3D – no-substrate 3D culture control, SM – smooth, DIM – dimpled, HD – heterogeneously dimpled.*

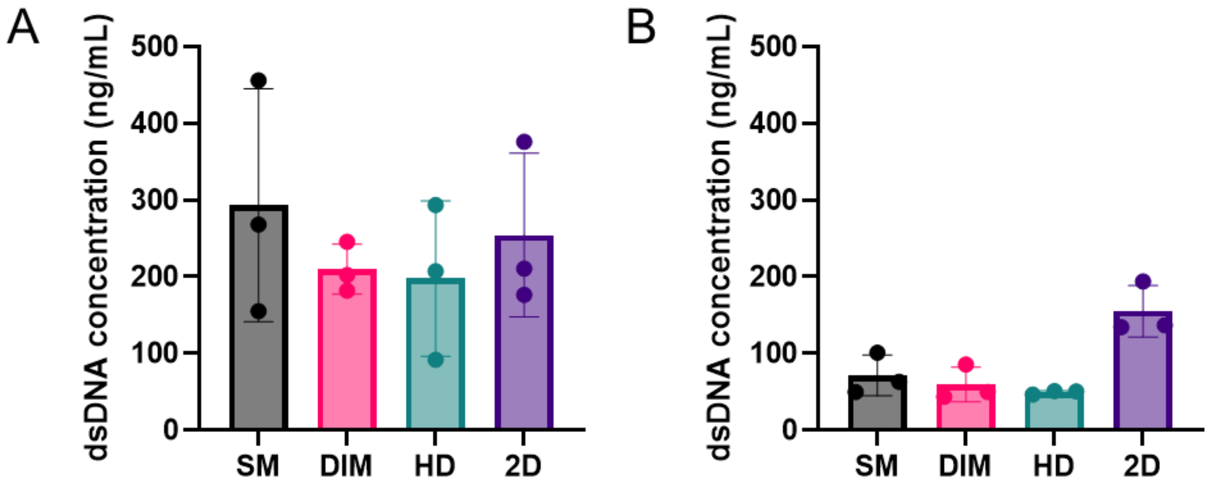


***Figure S6. Quantification of dsDNA content at 24 hours post-seeding.*** *PicoGreen assay was used to assess total dsDNA concentration of (A) MG-63 and (B) U2OS at 24 hours post seeding (n=3). Friedman test with Dunn's multiple comparisons test was performed.*
*Abbreviations: 2D – monolayer culture control, SM – smooth, DIM – dimpled, HD – heterogeneously dimpled.*

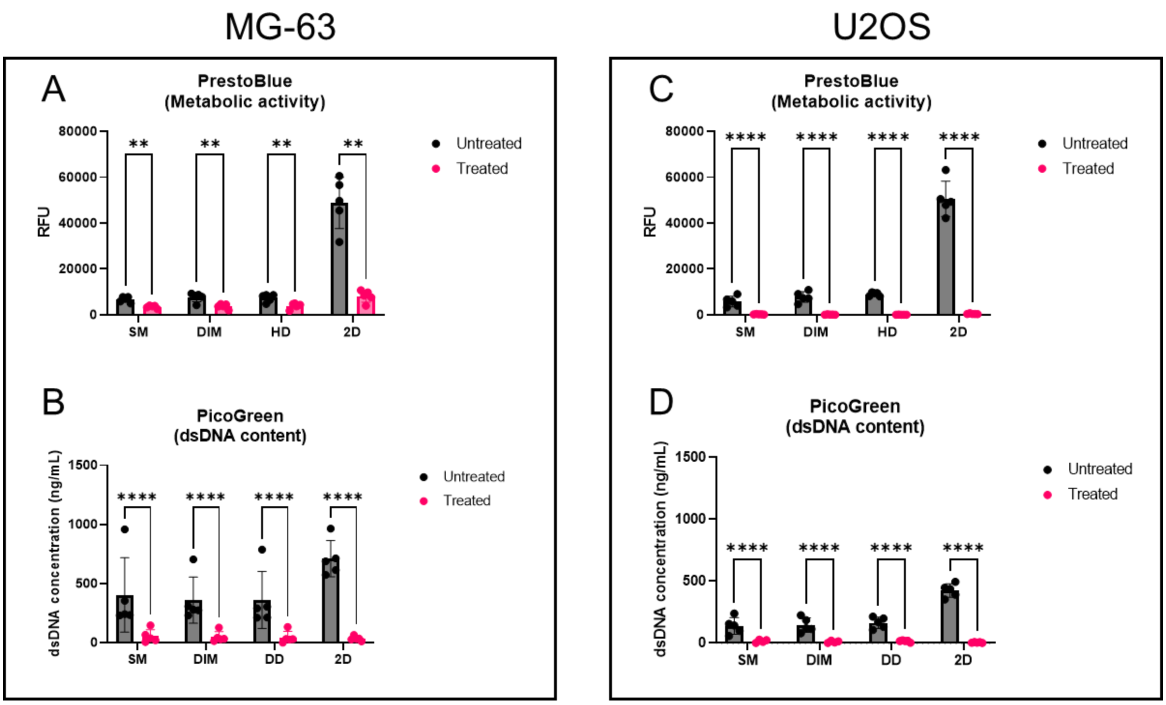


***Figure S7. Difference in metabolic activity and dsDNA content post doxorubicin treatment in MG-63 and U2OS cells.*** *Change in (A) metabolic activity and (B) dsDNA content after treatment with 10 µM doxorubicin in MG-63 cells. Change in (C) metabolic activity and (D) dsDNA content after treatment with 10 µM doxorubicin in U2OS cells (n=4-5). Ordinary two-way ANOVA with Tukey's multiple comparisons test was performed. **-p<0.01, ****-p<0.0001*

*Abbreviations: 2D – monolayer culture control, SM – smooth, DIM – dimpled, HD – heterogeneously dimpled.*

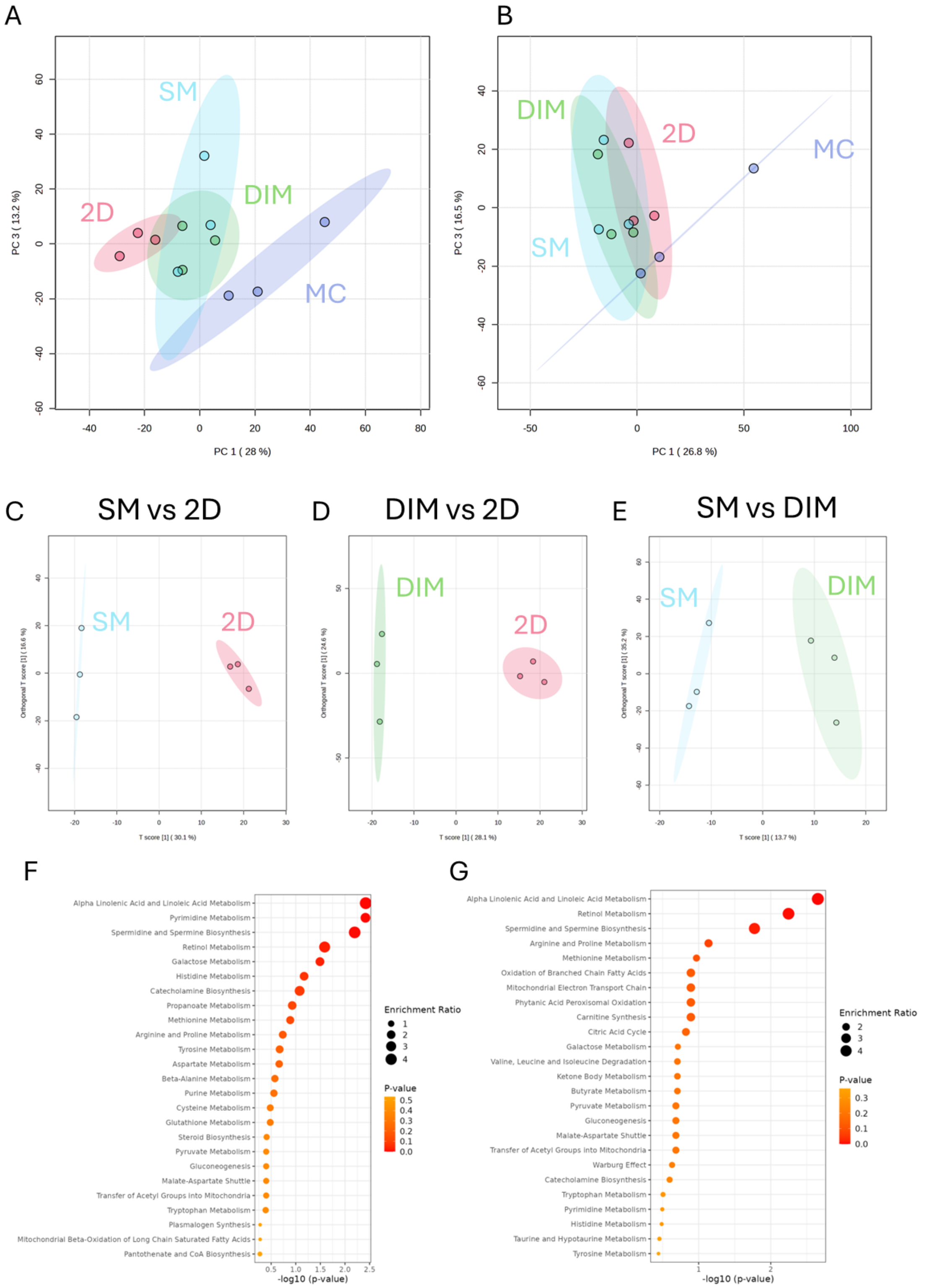


***Figure S8. Multivariate and enrichment analysis of metabolic profiles from MG-63 cultures.*** *(A, B) Principal component analysis (PCA) plots of metabolic profiles for (A) untreated and (B) treated cultures. (C-E) Orthogonal PLS-DA plots for doxorubicin-treated samples showing comparisons of (C) smooth microparticle cultures versus 2D, (D) dimpled microparticle versus 2D cultures, and (E) smooth versus dimpled microparticle cultures. (F, G) Top 25 enriched metabolite sets shown for doxorubicin-treated samples (F) smooth vs 2D cultures and (G) dimpled vs 2D cultures. PLS-DA, oPLS-DA and quantitative enrichment analyses performed on MetaboAnalyst 6.0. Abbreviations: SM – smooth, DIM – dimpled, MC – media control.*

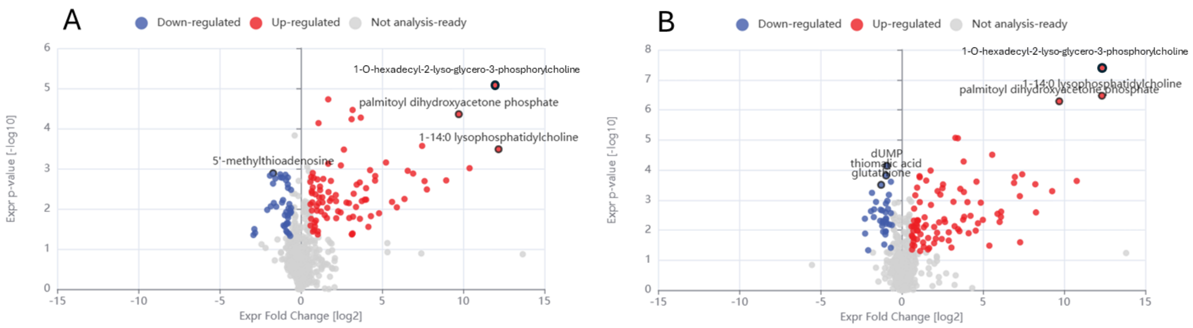


***Figure S9. Volcano plots showing differentially abundant extracellular metabolites in MG-63 cultures ($p < 0.05$, $|FC| > 1.5$).*** *(A) Differentially downregulated and upregulated metabolites in smooth vs 2D cultures. (B) Differentially down- and upregulated metabolites in dimpled microparticle vs 2D cultures.*

*Table S1. Predicted upstream regulators and toxicological functions altered in smooth vs 2D cultures (p < 0.05), based on Ingenuity Pathway Analysis. Directionality is based on activation z-scores, indicating predicted upregulation or downregulation.*

| **Upstream Regulators** | **P-value** | **Activation z-score** |
|---|---|---|
| ChoK | 0.00308 | 1.72 |
| LDL | 0.01 | 0.29 |
| chlorpyrifos | 0.01 | 1.01 |
| 3-nitropropionic acid | 0.01 | 1.07 |
| eicosapentaenoic acid | 0.01 | 0.15 |
| L-arginine | 0.01 | -0.83 |
| EGF | 0.02 | -0.82 |
| N-propyl bromide | 0.02 | -1.09 |
| GAC0001E5 | 0.03 | 1.13 |
| **Toxicological functions** | **P-value** | **Activation z-score** |
| Increased activation of alkaline phosphatase | 0.000584 | 1.21 |

Abbreviations: ChoK – choline kinase alpha, LDL – low density lipoprotein, EGF – epidermal growth factor

*Table S2. Predicted upstream regulators and toxicological functions altered in dimpled vs 2D cultures (p < 0.05), as identified by Ingenuity Pathway Analysis. Directionality is based on activation z-scores, indicating predicted upregulation or downregulation.*

| **Upstream Regulators** | ***p*-value** | **Activation z-score** |
|---|---|---|
| ChoK | 0.00516 | 1.72 |
| LDL | 0.02 | 0.29 |
| chlorpyrifos | 0.02 | 1.01 |
| eicosapentaenoic acid | 0.02 | 0.15 |
| EGF | 0.02 | -0.82 |
| **Toxicological functions** | ***p*-value** | **Activation z-score** |
| Increased activation of alkaline phosphatase | 0.0055 | 0.85 |

Abbreviations: ChoK – choline kinase alpha, LDL – low density lipoprotein, EGF – epidermal growth factor
